# Sociality in weevils is shaped by sheltering and convergent gene losses

**DOI:** 10.64898/2026.09.19.752859

**Authors:** Sarah Rinke, James Bickerstaff, Peter Biedermann, Carsten Kemena-Rinke, Markus Riegler, Martin Schebeck, Mark C. Harrison

## Abstract

Eusociality, characterized by overlapping generations, cooperative brood care, and reproductive division of labour, has arisen independently across diverse, phylogenetically distant insect orders, including Hymenoptera (ants, bees, and wasps), Blattodea (termites), and Coleoptera (weevils). While multiple studies have investigated the molecular evolution of sociality from solitary ancestors in Hymenoptera and Blattodea, so far little is known about the evolutionary signatures of social evolution in Coleoptera. Weevils (Curculionidae) provide an ideal system for addressing this question, as they cover the full spectrum of social complexity from parental care, through several origins of facultative eusociality to the only obligately eusocial beetle, *Austroplatypus incompertus*. We generated genome assemblies for *A. incompertus* and two facultatively eusocial weevil species, *Xylosandrus germanus* and *Xyleborinus saxesenii*, which together with 18 publicly available weevil genomes span two independent evolutionary origins of sociality. Our analyses reveal a genome-wide relaxation of purifying selection with increasing social complexity, which is most pronounced in *A. incompertus*. We find a significant excess of convergent gene family contractions in lineages where sociality evolved, and no evidence of elevated positive selection. These findings indicate that the molecular mechanisms of social evolution in weevils are primarily characterised by relaxed selection and gene loss, rather than adaptive innovation and gene family expansions. These observations are consistent with sheltering and reduced effective population size playing an important role, a pattern not previously observed in other clades.

**Significance statement:** Eusociality has evolved independently multiple times across the insect tree of life, yet the genomic mechanisms underlying these transitions remain poorly understood outside of Hymenoptera and termites. By generating genomes for *Austroplatypus incompertus*, the sole obligately eusocial beetle, and closely related facultatively eusocial weevil species, we construct a comprehensive genomic dataset spanning the full spectrum of social complexity within a single family. Genome-wide relaxation of purifying selection accompanies increasing social complexity in weevils, consistent with patterns documented in other social insects. Interestingly, instead of positive selection and gene family expansions, the main genomic signature is one of convergent gene family contractions. These findings demonstrate that the genomic routes to eusociality are more diverse than previously appreciated.

## Introduction

Insects exhibit a continuum of social behaviours, from solitary living to the most complex levels of eusociality or superorganismality (Boomsma & Gawne 2018). Eusociality is classically defined by cooperative brood care, overlapping generations, where offspring stay in the nest and often help their parents, and division of reproductive labour between reproductives and sterile workers (Wilson et al. 1971). Besides giving rise to ant, bee and wasp societies within Hymenoptera, eusociality along with various other social forms has evolved within Blattodea (termites), beetles, aphids and thrips (Costa 2006).

In beetles, different levels of social behaviour have evolved in at least ten families, in some even multiple times independently (Biedermann & Nuotclà 2020; Costa 2006). These diverse social behaviours within beetles are often linked to the use of ephemeral resources such as decaying plant material, dung or carcasses (Kölliker 2012). Cooperation between a male and a female beetle and parental care evolved to defend these resources from competitors, and process them for their offspring. For example, in burying beetles, rapid hiding and preparation of the carcass by both parents ensures sufficient resources for their larvae (Eggert & Müller 1997). Maternal care of offspring (e.g. protection, grooming and feeding of larvae) has evolved in Chrysomelidae, Erotylidae, Lucanidae and Staphylinidae, while biparental care has evolved in Scarabaeidae, Silphidae, and possibly Silvanidae. Alloparental care, i.e. helpers providing care to brood that are not their own, has evolved in Passalidae, Tenebrionidae and Curculionidae (Biedermann & Nuotclà 2020). Within Curculionidae, reproductive division of labour also evolved in one species, the pinhole borer *Austroplatypus incompertus*, making it the only eusocial beetle (Kent & Simpson 1992; Smith et al. 2018).

The entire spectrum of insect sociality exists within Curculionidae, the true weevils. While most weevils have a solitary lifestyle, high social complexity evolved within fungus-farming weevils (ambrosia beetles). Although the term “ambrosia beetles” may also refer to beetles outside of weevils, in this work we use it only for fungus-farming weevils, which encompass all Platypodinae and several clades of Scolytinae (Kirkendall et al. 2015). Parental care evolved multiple times within weevils, mainly within the wood-boring bark beetles (Scolytinae) and pinhole borers (Platypodinae) (Biedermann & Nuotclà 2020). Many weevil species, in which parental care is performed, live in tunnels within living or decaying trees (Kirkendall et al. 2015). Bark beetles (Scolytinae) excavate galleries mainly in the phloem and feed directly on tree tissues, while associated fungi are generally facultative mutualists (Six 2012). As bark beetles get most of their nutrients directly from their host plant, they tend to be more specialised on certain tree taxa. Many bark beetles are at most subsocial, aggregating to perform mass attacks on trees and performing some parental care behaviour like egg and larval attendance (Blomquist et al. 2010; Kirkendall et al. 2015). Additional social behaviour, like alloparental care and overlapping generations, is especially prevalent in fungus-farming ambrosia beetles. These beetles bore into nutrient-poor xylem and rely on their obligate symbiotic fungi for nutrients (Six 2012). The beetles evolved a mycetangium to bring fungal spores to new nests and inoculate the gallery walls to farm fungi as their only food source. Often, ambrosia beetles have a broader host tree range as their associated fungi enable them to acquire nutrients from different host plants (Hulcr & Stelinski 2017). This labour-intensive feeding behaviour favours the evolution of cooperative brood care as well as reproductive division of labour (Biedermann & Rohlfs 2017). In some ambrosia beetles, for example *Xyleborus* and *Xylosandrus* species (Scolytinae), female offspring delay their dispersal from the parental nest in order to support the colony in the cultivation of fungus and the rearing of siblings, while still retaining their ability to reproduce (Biedermann et al. 2011; Kirkendall et al. 2015). Large groups of beetles (typically 20, but >100 are possible) living in narrow tunnel systems are expected to select for social immunity, such as removing frass and grooming larvae, as well as collective pathogen defences. As galleries and associated fungi are transient but defensible, group living and (allo-)parental care increase food availability, safety and offspring survival, leading to multiple independent origins of social behaviour in these beetles (Biedermann & Rohlfs 2017).

Besides exhibiting eusociality, *A. incompertus* differs from other weevils in several ecological and reproductive traits. This species is distributed in discontinuous patches of mesic forests along the eastern Australian coastline where it constructs nests in healthy living stringybark *Eucalyptus* hosts (Bickerstaff et al. 2020; Kent 2008). Despite substantial genomic divergence throughout the species’ range (Bickerstaff et al. 2025), all populations are consistently associated with a single *Raffaelea* symbiotic fungus (Mueller 2019). Nest construction and fungus activity has minimal impact on overall host-tree health, likely contributing to the longevity (up to 36 years) of the species and gallery systems (Harris et al. 1976; Kent & Simpson 1992; Smith et al. 2018). Colonies typically comprise a single inseminated foundress, her unmated daughter workers, and immature developmental stages of both sexes (Smith et al. 2018). All adult males and a subset of adult females disperse from the colony to mate with unrelated individuals in somewhat close (< 500m) proximity to their natal nest (Smith 2013). Despite their low dispersal capacity, and unlike social Xyleborini, colonies are characterised by outbreeding and are diplodiploid (Bickerstaff et al. 2025; Smith et al. 2009). No external morphological caste differentiation exists between female caste members, with only non-reproductive daughter workers having empty spermathecae and vestigial ovaries (Smith et al. 2018).

Social traits are expected to influence genome evolution, leaving measurable traces in gene family sizes, selection patterns, and molecular evolutionary rates. Colony size and generation time, two important characteristics of social complexity in insects, have been shown to significantly correlate positively and negatively, respectively, with molecular evolutionary rate in Hymenoptera (Rubin 2022). The relationship between genome content and social complexity is unclear. Within Hymenoptera eusocial species tend to possess larger genomes than solitary species, while the opposite is true for Blattodea (Mikhailova et al. 2024). Furthermore, transposable element (TE) content is negatively correlated with social complexity in bees (Kapheim et al. 2015), positively correlated with sociality in snapping shrimps (Chak et al. 2021), while no relationship is apparent in Blattodea (Mikhailova et al. 2024). Multiple molecular mechanisms have been implicated in the evolution of social complexity. Transcriptional regulation appears relevant across social complexity levels: changes in the regulation of gene expression are predicted to be an important mechanism in the evolution of subsociality, and caste-biased gene expression underpins worker-reproductive divergence in eusocial species (Mikhailova et al. 2024; Rehan & Toth 2015). High numbers of genes under positive selection and an increased importance of clade-specific genes are also associated with transitions to higher social complexity levels (Rehan & Toth 2015).

Relaxed purifying selection has repeatedly been associated with a decrease in effective population size due to reproductive skew in eusocial colonies, both in Hymenoptera (Weyna & Romiguier 2021) and termites (Ewart et al. 2024; Roux et al. 2024). However, further studies indicate a reduction in selection strength may predate the evolution of sociality in these lineages (Hunt et al. 2011; Jones et al. 2026). In fact, parental care of offspring may also relax selection due to a sheltering effect, as shown in the subsocial carrion beetle *Nicrophorus vespilloides* (Mashoodh et al. 2023). In support, the first study addressing sociality in weevils found that relaxation of selection and convergent gene loss are associated with the emergence of parental care (Rinke et al. 2026).

We have compiled a comprehensive genomic dataset comprising 18 high-quality, publicly available Curculionidae genomes, which we supplement with three new genome assemblies, including that of the only eusocial beetle, *A. incompertus*. Here, we use this dataset to investigate the molecular mechanisms involved in the evolution of sociality across various weevil families. We hypothesised that the evolution of sociality in beetles is accompanied by similar mechanisms already observed in other clades, i.e. transcriptional regulation and relaxation of selection playing a role in the evolution of lower social complexity levels, while positive selection and clade specific genes may play a role in the evolution of more complex sociality.

## Results

### Large dataset of high quality weevil genomes, covering a broad range of social traits

Our dataset comprises genomes of 21 Curculionidae species, with the leaf-rolling weevil, *Apoderus coryli* (Attelabidae), and the red flour beetle,*Tribolium castaneaum* (Tenebrionidae), as outgroups. With these genomes, we cover a broad range of combinations of the social traits nest building, egg attendance, larval attendance, overlapping generations, alloparental care, and sterile workers (Fig. 1). All 20 publicly available genome assemblies used in this study are of high quality, according to both, assembly (N50 *>* 125kb, completeness *>*95%) and annotation statistics (completeness *>*90%; Table S2; Fig. 2). These were complemented by the newly sequenced genomes of the two highly social ambrosia beetles *Xylosandrus germanus*, *Xyleborinus saxesenii* and the only eusocial beetle, *A. incompertus* (Fig. 1, species in bold font). We produced high quality assemblies for *X. germanus* (N50 = 16.3 Mb and L50 = 5) and *X. saxesenii* (N50 = 10.8 Mb and L50 = 7) using PacBio Hifi. The genome assembly of *A. incompertus* is less contiguous (N50 = 194 kb and L50 = 140), due to reduced sample quality and different sequencing and assembly methods (ONT PromethION and Illumina). All three new genomes are highly complete (assemblies: *>* 96%; annotations: *>* 94%; see Fig. 2 and Table S2).

**Figure 1:**
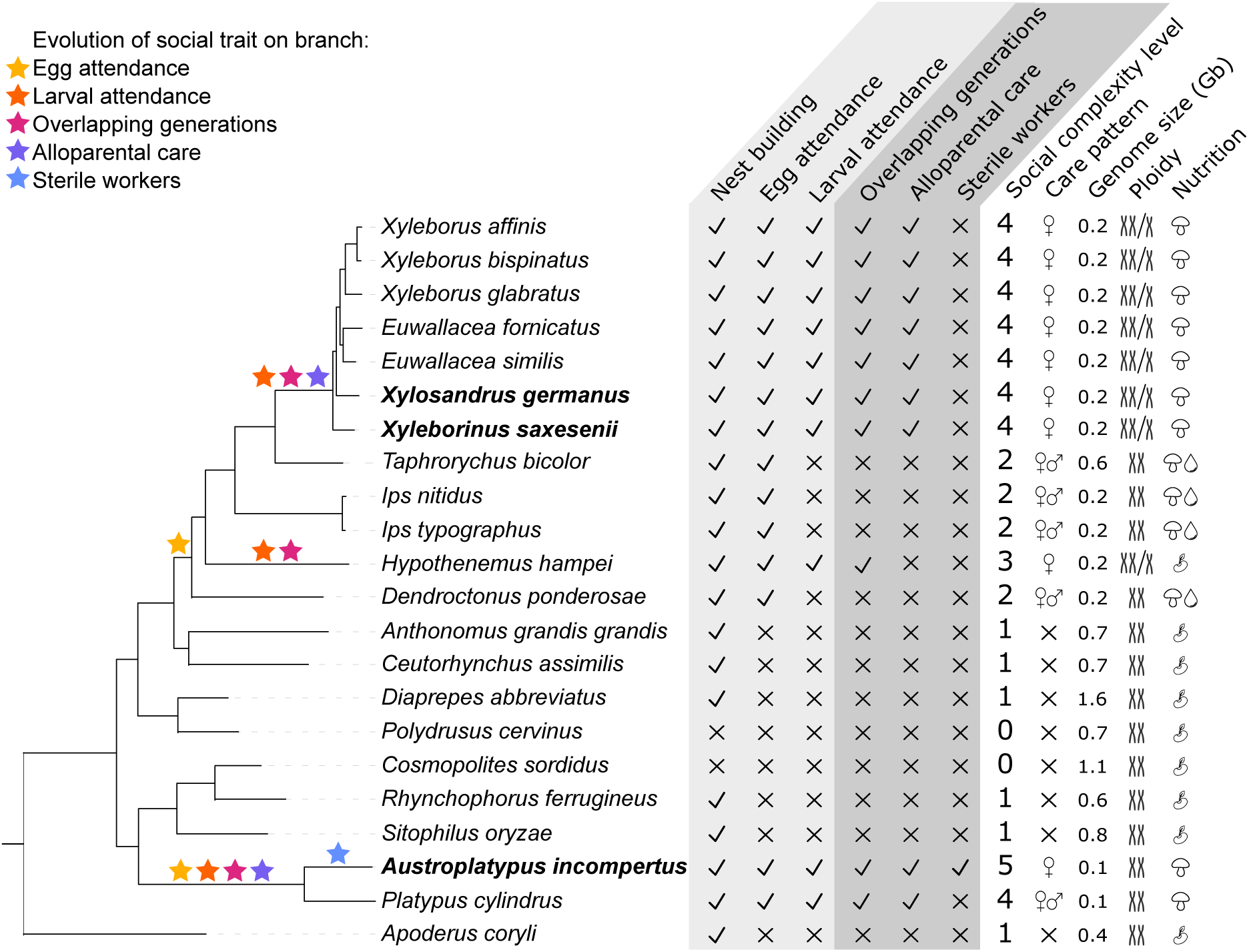
Phylogenetic tree of species used in this study with social traits and other phenotypes. Stars indicate the branch where each social trait evolved. These social traits are also shown for each species in the table next to the phylogenetic tree, where tick marks mean presence of trait and x means absence of trait. Social complexity level is an ordinal representation of which social traits are present in each species: 0 = no social trait, 1 = nest building only, 2 = nest building + egg attendance, 3 = nest building + egg attendance + larval attendance + overlapping generations, 4 = nest building + egg attendance + larval attendance + overlapping generations + alloparental care, and 5 = nest building + egg attendance + larval attendance + overlapping generations + alloparental care + sterile workers. Additionally, care pattern (none, female or biparental), genome size in gigabases, ploidy (diploid or haplo-diploid) and nutrition (plant parts, fungi, fungi and phloem) are shown. Species names in bold indicate genomes sequenced and assembled in this study.

**Figure 2:**
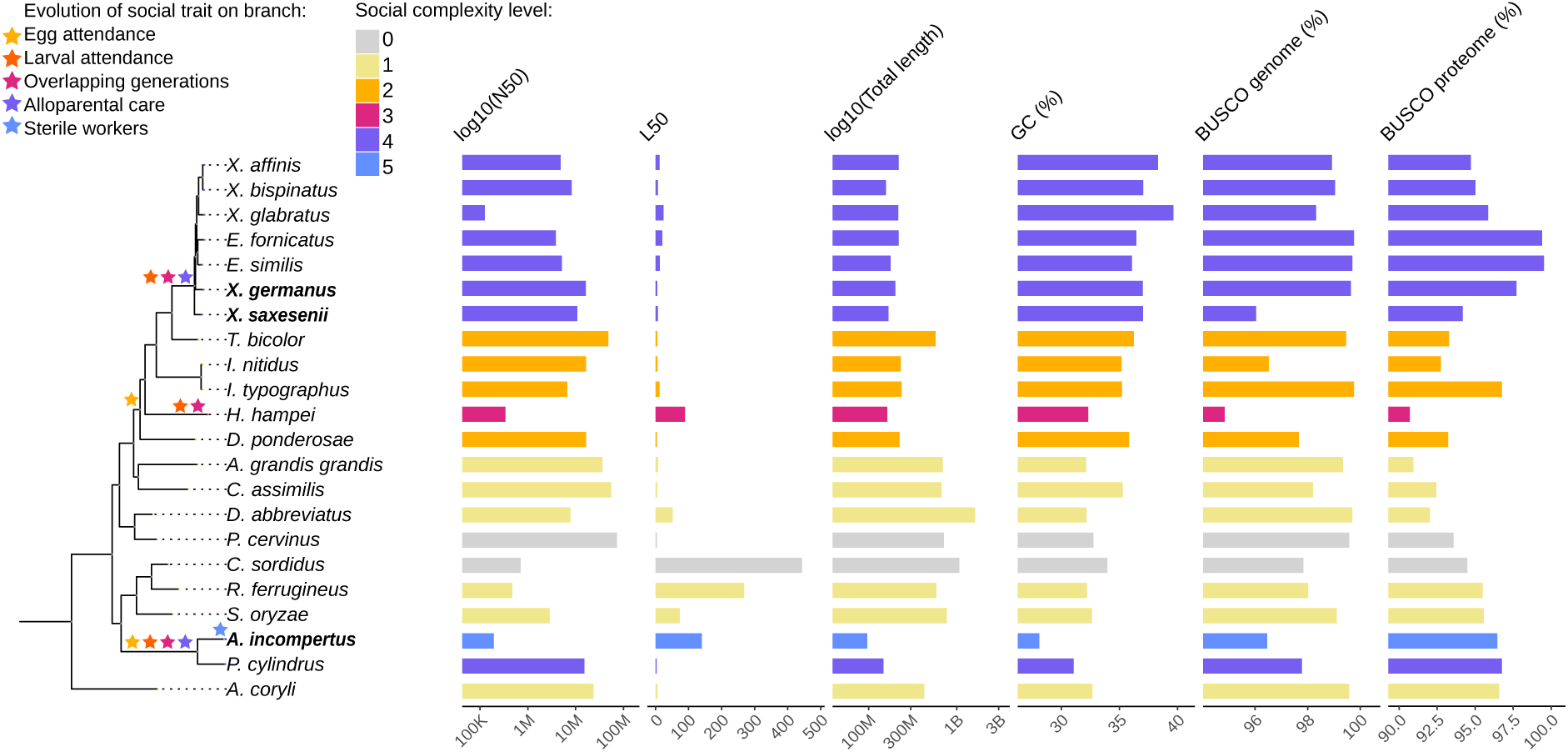
Overview of assembly and annotation statistics. For each species, N50, L50, total length, GC content, BUSCO completeness score for genomes and proteomes are shown as barplots. Colours of bars indicate social complexity level, an ordinal representation of which social traits are present in each species: 0 = no social trait, 1 = nestbuilding only, 2 = nest building + egg attendance, 3 = nest building + egg attendance + larval attendance + overlapping generations, 4 = nest building + egg attendance + larval attendance + overlapping generations + alloparental care, and 5 = nest building + egg attendance + larval attendance + overlapping generations + alloparental care + sterile workers. Stars in the tree indicate evolutionary origins of social traits.

The genome assembly lengths of these three highly social species is below the mean (427.8 Mb) for the dataset, with the eusocial *A. incompertus* having the shortest genome of all (*∼* 96.4 Mb, see Fig. 2, Table S2). Repetitive content and genome size have a strong positive correlation (r*_s_* = 0.74, p = 0.0001; Fig. S1) and both also have a strong negative correlation with sociality level, Spearman’s rank correlation r = -0.795 and r = -0.861, respectively (Fig. S2). However, a Phylogenetic Generalized Least Squares analysis (PGLS) showed no significance for the correlation between sociality and genome size (p = 0.18) or for the correlation between sociality and repetitive content (p=0.22) when taking phylogeny into account. GC content varies greatly across the species in this dataset (28.11% in *A. incompertus* -39.65% in *Xyleborus glabratus*, see Fig. 2, Table S2), with all the Xyleborini species having higher GC content than all other species. After correcting for phylogenetic relationship, GC content is not correlated with social complexity level (p=0.56). Total number of annotated protein coding genes varied between 10,663 in *Euwallacea fornicatus* and 41,176 in *X. glabratus*, with most species having fewer than 20,000 protein coding genes (Table 1). The obligately eusocial *A. incompertus* has the second lowest number of protein coding genes, namely 12,034 but there is no significant correlation p=0.99) between number of protein coding genes and social complexity level after correcting for phylogeny.

**Table 1:**
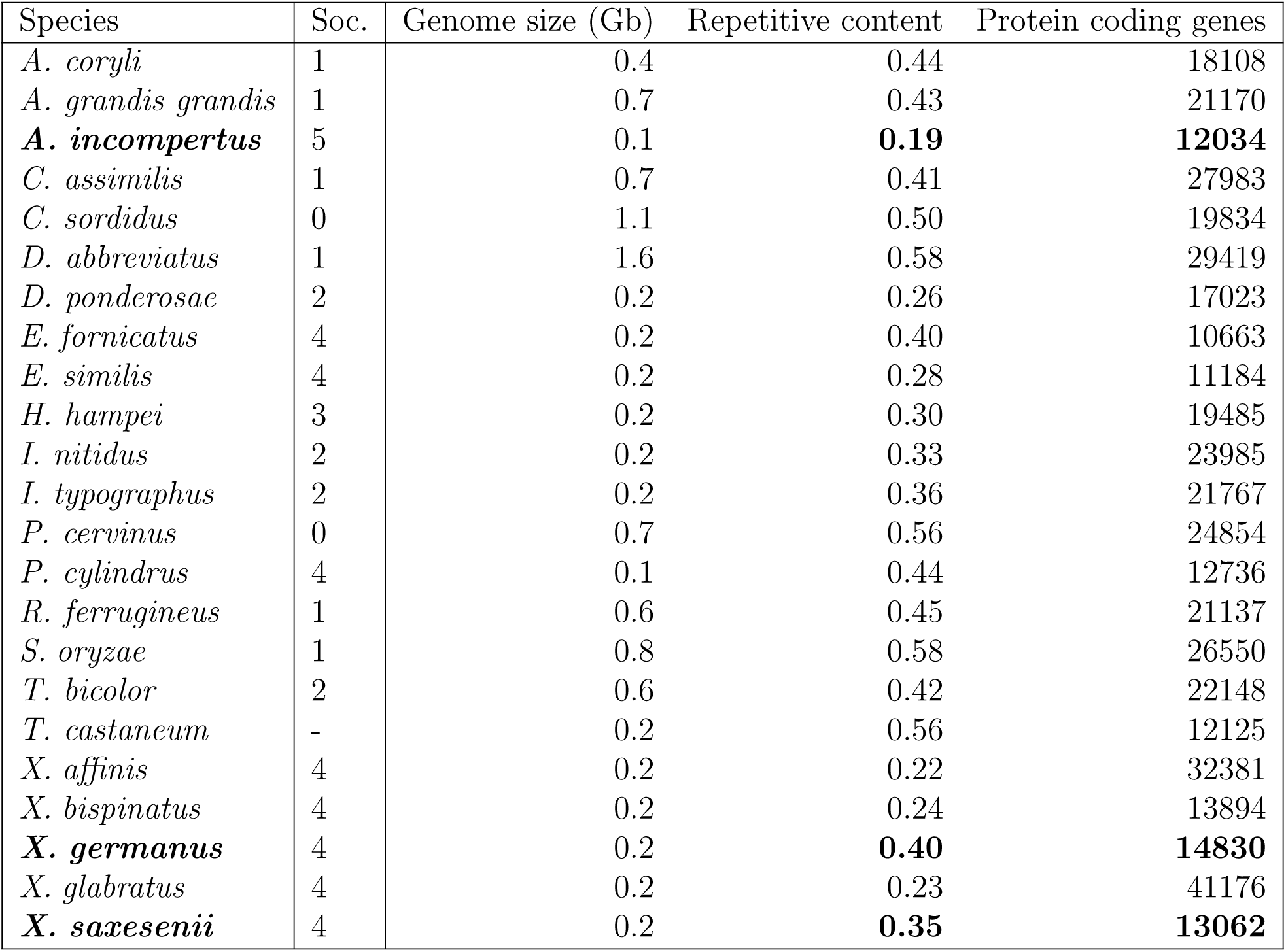
Repetitive genome content and numbers of protein coding genes. Percentage of repetitive content was calculated from soft-masked genomes. Number of protein coding genes was extracted from proteomes. Soc. = Social complexity, see Fig. 1 for details on social complexity levels.

### Weevils span the entire continuum of sociality and have multiple independent origins of social traits

The analysed weevil genomes cover multiple subfamilies, including Scolytinae and Platypodinae, where sociality evolved. Within the Platypodinae, only *A. incompertus* (Fig. 3 c, assembled in this study) and *Platypus cylindrus* (Barclay et al. 2024) are included; the former is eusocial and the latter is highly social but lacking a permanently sterile worker caste. Within the Scolytinae, twelve species were included. Seven species belong to the Xyleborini tribe and all exhibit similar social traits to *P. cylindrus*, performing parental care and alloparental care, as well as having overlapping generations. The species included from this subfamily are *Xyleborus affinis* (Iridian Genomes), *Xyleborus bispinatus* (Iridian Genomes), *X. glabratus* (Iridian Genomes), *E. fornicatus* (Bickerstaff et al. 2024), *Euwallacea similis* (Bickerstaff et al. 2024), *X. germanus* (Fig. 3 a, assembled in this study) and *X. saxesenii* (Fig. 3 b, assembled in this study). A further weevil species within this dataset that exhibits greater social complexity than just egg attendance is *Hypothenemus hampei* (Navarro-Escalante et al. 2021), which performs larval attendance and has overlapping generations. The other Scolytinae species included in this study all perform egg attendance: *Taphrorychus bicolor* (Telfer & Badham 2024), *Ips nitidus* (Wang et al. 2023), *Ips typographus* (Powell et al. 2021) and *Dendroctonus ponderosae* (Keeling et al. 2022). Five weevils from other families included in this study only build nests: *Anthonomus grandis grandis* (Childers et al. 2021), *Ceutorhynchus assimilis* (Pest Genomics Initiative between Rothamsted Research, Bayer, and Syngenta), *Diaprepes abbreviatus* (Sylvester et al. 2024), *Rhynchophorus ferrugineus* (Dias et al. 2021) and *Sitophilus oryzae* (Parisot et al. 2021); while *Polydrusus cervinus* (Barclay et al. 2023) and *Cosmopolites sordidus* (Rodriguez Ruiz & Van Dam 2023) do not exhibit care traits (Fig. 1).

**Figure 3:**
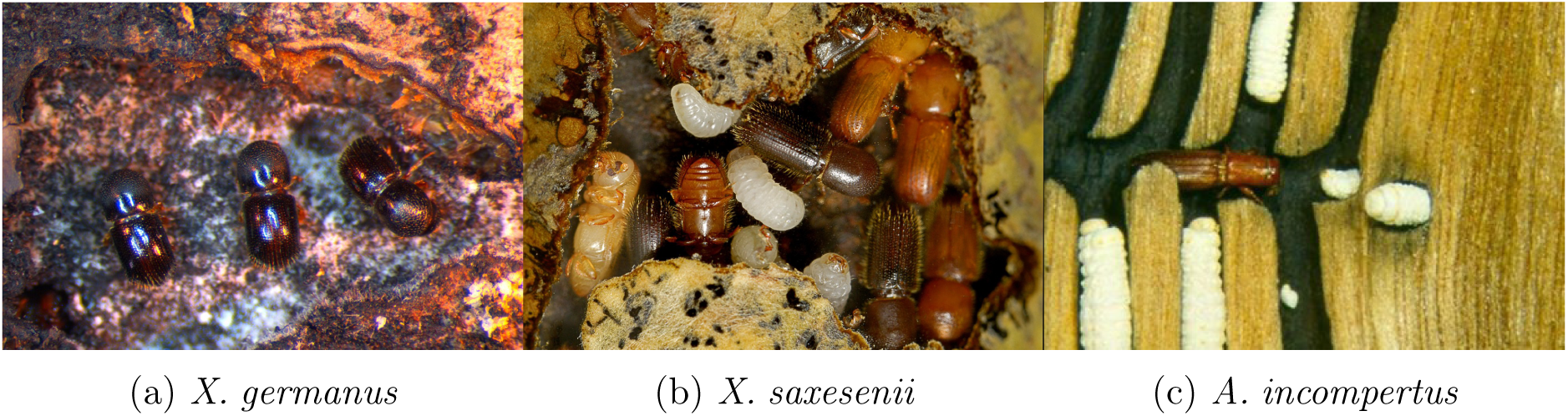
Photos of social weevils sequenced in this project. a) Brood chamber of *X. germanus* within artificial sawdust medium. Mature females are jointly tending the nutritional fungus, which produces white “ambrosial growth” structures along the walls of the chamber. Photo credit: Christopher Ranger. b) Brood chamber of *X. saxesenii* within artificial sawdust medium. Second- and third-instar larvae, teneral (light brown), and mature (black) females are jointly tending each other and the fungus. Yellow fungal hyphae of the nutritional fungus are growing into the medium and excrete melanin (black spots). Photo credit: Davide Vallotto. c) Gallery of adult and larval *A. incompertus* with dark fungal staining of the gallery. Photo credit: Geoff Avern.

To determine where in the phylogeny social traits evolved, we performed ancestral reconstruction for nest building, egg attendance, larval attendance, overlapping generations, alloparental care and sterile workers. The evolution of nest building is least clear within this dataset (Fig. S3 a) and its evolution is not further analysed (similarly to Rinke et al. 2026). Egg attendance evolved twice within this dataset, once at the root of all Platypodinae and once at the root of all Scolytinae (see Fig. S3 b, yellow stars in Fig. 1). Larval attendance and overlapping generations evolved on the same branches within this dataset, once in *H. hampei* and once in the Platypodinae ancestor (see Fig. S3 c and d, orange and pink stars in Fig. 1). Alloparental care evolved in the Platypodinae ancestor and Xyleborini ancestor (see Fig. S3 e, purple stars in Fig. 1). Sterile workers, one of the hallmarks of eusociality, only evolved in *A. incompertus* (see Fig. S3 f, blue star in Fig. 1).

### Low substitution rates in species with alloparental care

We measured variation in synonymous and non-synonymous substitution rates (dN and dS, respectively, see Fig. S5 and Fig. S6) to infer potential effects of social traits on mutation rates. As expected for social species with longer generation times, we found substantially lower dN and dS in social species that perform alloparental care (mean dN: 0.009; mean dS: 0.049) compared to those that do not (mean dN: 0.038; mean dS: 0.210; FDR: 0.007 & 0.005). A PGLS revealed a large standardised effect size of -0.928 and -0.916 (SD units, phylogenetically corrected) for dN and dS, respectively, when comparing species with and without alloparental care, despite a strong phylogenetic signal (Pagel’s *λ*: 1.0). The non-significant PGLS analysis was therefore likely caused by a limited number of independent origins (two) of alloparental care in our dataset. Egg attendance (−0.48) and larval attendance (−0.33 & -0.31), on the other hand, had only a small to moderate standardised effect on dN and dS, when controlling for phylogeny. Overall, these results confirm the expectation that socially complex species experience reduced mutation rates due to longer generation times than less social species.

### Contraction of diverse gene families at the origins of social traits

We used CAFE5 (Mendes et al. 2021) to analyse how patterns of gene family size evolution relate to the emergence of social traits in our dataset. Generally, there are more expansions and contractions on the terminal than on ancestral branches, but we found no obvious pattern at the branches of interest (stars in Fig. S7).

Comparing the significantly expanded and contracted gene families on all branches of interest, *A. incompertus* and *H. hampei* have more expansions than contractions, and approximately tenfold more expansions than the ancestral branches of interest (ancestors of Xyleborini, Scolytinae and Platypodinae; Table S3). On these three ancestral branches of interest, there are more contractions than expansions.

GO term enrichment analysis showed no functional enrichment within the gene families that were significantly expanded on any ancestral branch of interest (Scolytinae, Platypodinae, and Xyleborini ancestors). However, on the *H. hampei* branch, where larval attendance and overlapping generations evolved, significantly expanded gene families were enriched for the GO terms signal transduction, cell communication, cellular response to stimulus, response to stimulus, signaling and phosphorus metabolic process. Significant expansions on the *A. incompertus* branch, where sterile workers evolved, were enriched for the term microtubule-based process. None of these terms was significant after FDR correction (FDR *≥* 0.05; Table S3).

The gene families that were contracted on the *A. incompertus* branch, were significantly enriched for the functions response to stress and DNA metabolic process (FDR *<* 0.05), while the GO-terms regulation of molecular function, multicellular organismal process, cellular response to stress and DNA repair were only significant prior to FDR correction (p *<* 0.05). In *H. hampei*, significantly contracted gene families were enriched for proteolysis, protein metabolic process, aromatic compound biosynthetic process, organic cyclic compound biosynthetic process and heterocycle biosynthetic process prior to FDR correction (p *<* 0.05). Within contracted genes families in the Platypodinae ancestor, where multiple social traits emerged, the term multicellular organismal process was enriched significantly at an FDR corrected level, while transmembrane transport and proteolysis were only significant prior to FDR correction. In contracted gene families in the Scolytinae ancestor, where egg attendance evolved, only two GO terms were significantly enriched prior to FDR correction: DNA replication and proteolysis. The term proteolysis is the only term significantly enriched after FDR correction within contracted gene families in the Xyleborini ancestor (origin of larval attendance, overlapping generations, and alloparental care), while multicellular organismal process, cellular aromatic compound metabolic process, heterocycle metabolic process and DNA metabolic processes are significant before FDR correction. Interestingly, the term proteolysis is enriched in contractions on all branches of interest, except *A. incompertus*, while other terms, such as multicellular organismal process and DNA metabolic process, are enriched on multiple branches of interest (Table S3).

### Increased convergent gene losses along with evolution of sociality

To test for convergent gene family changes at the evolutionary origins of common social traits, we performed a SuperExactTest to see which branches had the largest and most significant overlaps in expanded or contracted gene families (Fig. 4). We found no significant convergent expansions between combinations of branches where social traits evolved. However, we found multiple occurrences of significant convergent contractions. At the two origins of egg attendance (the ancestral branches of Scolytinae and Platypodinae) eight gene families were convergently contracted (p = 3.47E-04; Fig. 4). At the three convergent origins of larval attendance and overlapping generations (Xyleborini ancestor, Platypodinae ancestor and *H. hampei*) six gene families were convergently contracted (p = 2.40E-10), while 15 were convergently contracted on the Platypodinae and *H. hampei* branches (p = 1.16E-11) and 11 shared between the Xyleborini and *H. hampei* origins of these two traits (p = 3.62E-06). On the Xyleborini and Platypodinae ancestral branches where beside larval attendance and overlapping generations, alloparental care also convergently evolved 19 gene families significantly contracted (p = 3.95E-09; Fig. 4).

**Figure 4:**
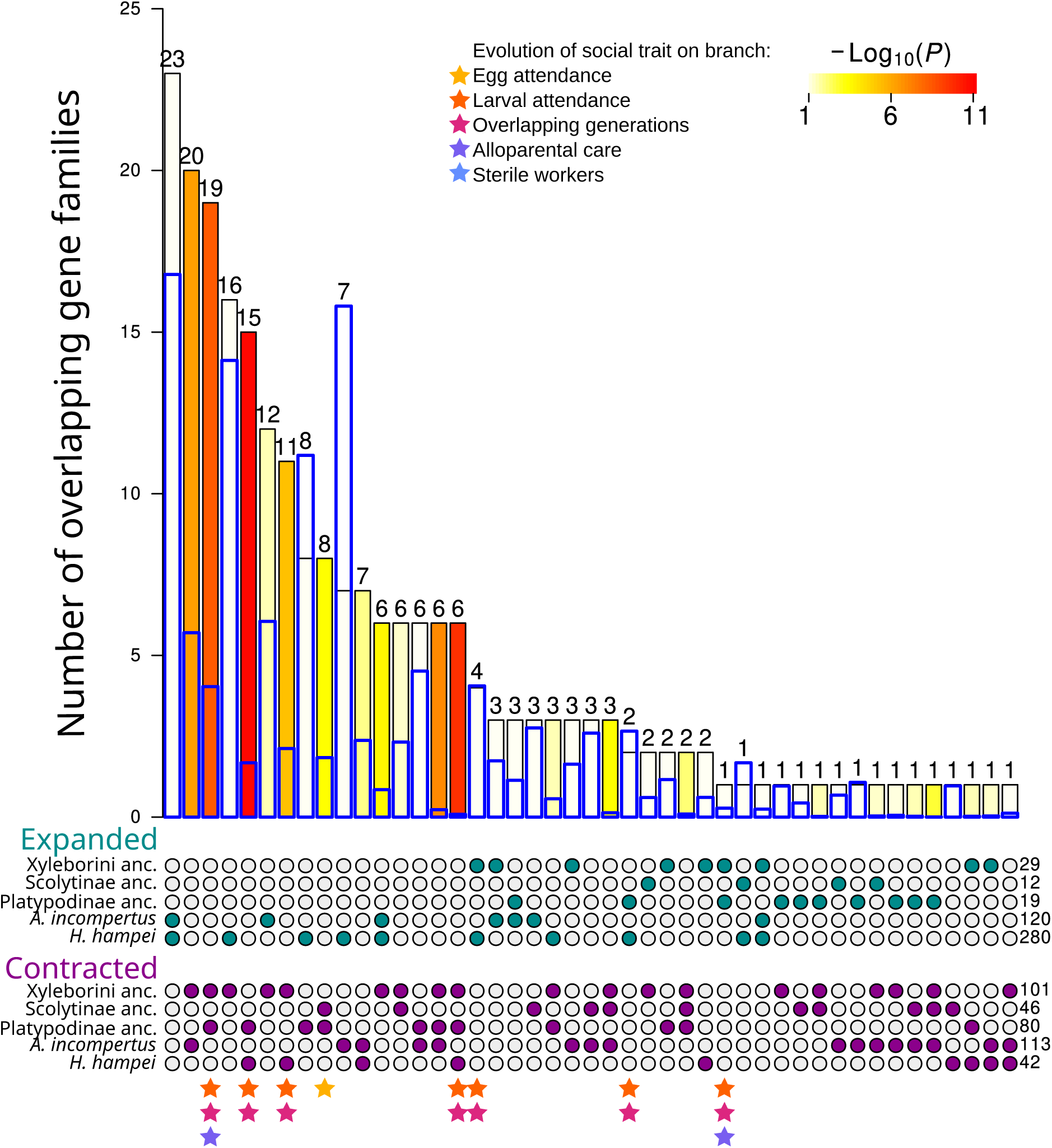
SuperExactTest of all overlaps of expanded and contracted gene families on all branches of interest. Turquoise dots (top five rows) correspond to expansions, while purple dots (bottom five rows) correspond to contractions. Blue thick boxes indicate the expected overlap, significance is shown in yellow-red scale as *−Log*_10_(*P*). Stars at the bottom indicate, which social traits have convergently evolved on the indicated branches: yellow (one star) -egg attendance, orange and pink (two stars) -larval attendance and overlapping generations, orange, pink and purple (three stars) -larval attendance, overlapping generations and alloparental care.

We investigated whether convergent gene family expansions and contractions were more common at origins of social traits than expected. For this we compared numbers of shared contractions and expansions between all pairs of branches within our phylogeny. On the whole convergent expansions of gene families on pairs of branches of interest did not differ from convergent expansions on other branch pairs (Mann-Whitney U=5999; p-value = 0.0721, Fig. 5). Interestingly, convergent contractions on two branches of interest were significantly overrepresented compared to convergent contractions on any two branches across the tree (Mann-Whitney U=2270; p-value = 0.0042, Fig. 5).

**Figure 5:**
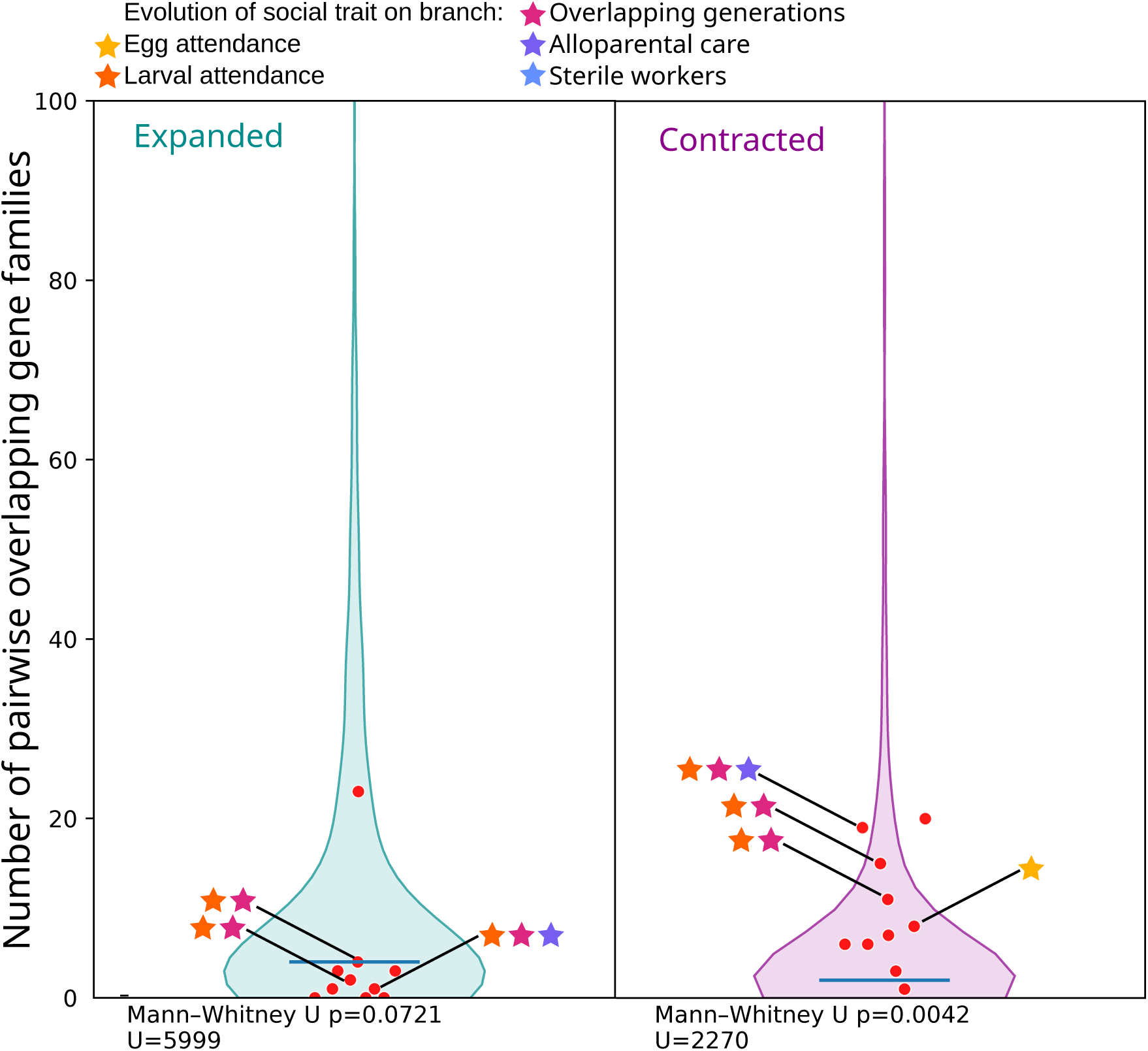
Distribution of all overlaps of expanded (left plot, turquoise) or contracted (right plot, purple) gene families between any two branches (violin plot) with the number of overlaps of contracted or expanded genes on each pair of branches of interest highlighted as a red dot. Number of combinations of two branches in the violin plots: contractions n=903, expansions n=946. Number of combinations of foreground branches: n=10. Stars indicate which social phenotype evolved on which pair of branches. yellow (one star) -egg attendance, orange and pink (two stars) -larval attendance and overlapping generations, orange, pink and purple (three stars) -larval attendance, overlapping generations and alloparental care.

The most interesting overlaps are the ones in the same direction (convergent contractions or expansions) on branches where the same social traits evolved. We will focus the description of functional annotations on these overlaps (Table S5. Functions associated with the 19 gene families convergently contracted on branches where larval attendance, overlapping generations and alloparental care evolved, namely on Xyleborini ancestor and Platypodinae ancestor, are very diverse and cover traits like sugar metabolism, stress response, lipid metabolism, detoxification, gene expression regulation and transport (Table 2).

**Table 2:** High-level functional annotation of convergently contracted and expanded gene families on different branches of interest where the same social phenotypes evolved. The overlaps correspond to overlaps between expansions or contractions marked with stars in the SuperExactTest (Fig. 4). The order corresponds to number of gene families per overlap (highest to lowest). Sociality indicates social phenotypes that evolved on these branches. Colours of stars and corresponding social phenotype: yellow (one star) -egg attendance, orange and pink (two stars) -larval attendance and overlapping generations, orange, pink and purple (three stars) -larval attendance, overlapping generations and alloparental care. Numbers indicate the count of gene families with annotations in each high-level functional category. All convergent expansion on branches where the same social phenotype evolved had non-significant results in the SuperExactTest, i.e. there were not more gene families convergently expanded than expected. More detail on the annotations of specific genes can be found in Table S5.

| Overlap | Sociality | Gene expression regulation <sup>a</sup> | Sugar metabolism <sup>b</sup> | Lipid metabolism <sup>c</sup> | Stress response <sup>d</sup> | Detoxification <sup>e</sup> | Transport <sup>f</sup> | Cuticular hydrocarbon synthesis <sup>g</sup> | Neuronal communication <sup>h</sup> | Others <sup>i</sup> | Not annotated |
| --- | --- | --- | --- | --- | --- | --- | --- | --- | --- | --- | --- |
| **Contractions** |  |  |  |  |  |  |  |  |  |  |  |
| Xyleborini ancestor & Platypodinae ancestor | ★ ★ ★ | 1 | 2 | 2 | 2 | 2 | 1 |  |  | 2 | 7 |
| Platypodinae ancestor & *H. hampei* | ★ ★ | 3 |  | 1 | 2 | 1 |  | 1 |  | 4 | 3 |
| Xyleborini ancestor & *H. hampei* | ★ ★ |  |  | 1 | 2 | 2 |  |  |  | 4 | 2 |
| Scolytinae ancestor & Platypodinae ancestor | ★ |  |  |  | 1 |  |  |  | 1 | 1 | 5 |
| Xyleborini ancestor & Platypodinae ancestor & *H. hampei* | ★ ★ |  |  | 1 | 2 |  |  |  |  | 2 | 1 |
| **Expansions** |  |  |  |  |  |  |  |  |  |  |  |
| Xyleborini ancestor & *H. hampei* | ★ ★ |  |  |  |  |  |  |  |  |  | 4 |
| Platypodinae ancestor & *H. hampei* | ★ ★ |  |  |  |  | 1 |  |  |  |  | 1 |
| Xyleborini ancestor & Platypodinae ancestor | ★ ★ ★ |  |  |  |  |  |  |  |  |  | 1 |

There are three sets of branches where larval attendance and overlapping generations evolved and which have significant convergent contractions in gene families, namely 15 gene families in Platypodinae ancestor and *H. hampei*, 11 gene families in *H. hampei* and the Xyleborini ancestor, and 6 in Xyleborini ancestor, Platypodinae ancestor and *H. hampei*. Although, each of these sets of gene families cover a large variety of functions, many of these functions are shared. Functional annotations for convergent contractions on all three branches where larval attendance and overlapping generations evolved (Xyleborini ancestor, Platypodinae ancestor and *H. hampei*) contain a “core set” of the above annotations: Heat shock protein, GMC-oxidoreductase, carboxyl-esterase, peptidase S1 and thrombospondin-type-1 (see Tables 2 and S5).

Additionally, there is one overlap of two branches where egg attendance evolved: in the ancestors of Scolytinae and Platypodinae are eight convergently contracted gene families. The functions for these convergently contracted genes are different, namely small heat shock protein, CRISP family and ionotropic glutamate-receptor activity. GO term enrichment analysis for this group revealed one significantly enriched term: multicellular organismal process.

### Developmental, immunity, and transcriptional regulatory gene families evolve specifically in *A. incompertus*

We identified 246 species-specific genes in *A. incompertus*. Although a substantial fraction of these genes lack functional annotation, those for which a putative function could be assigned span a diverse range of biological processes and can be broadly grouped according to the cellular or organismal traits they are likely to influence (Table S6).

Six genes can be associated with social biology and reproductive regulation, including a juvenile hormone-binding protein, two subunits of the heterotrimeric transcription factor NF-Y, a homeodomain-containing protein, as well as histone H1 and a histone acetylation factor. A further six genes are involved in signal transduction, predominantly encoding kinases and phosphatases such as two serine/threonine kinases, a tyrosine kinase, a phosphatidylinositol-specific phospholipase C, a myotubularin-family non-receptor protein tyrosine phosphatase, and a guanylate kinase homologue. Five genes are putatively linked to host-plant adaptation, most notably a cytochrome P450 implicated in detoxification, together with a trehalase, an enoyl-ACP reductase, an aspartate oxidase, and one oxidoreductase.

Four genes are related to protein turnover: an F-box protein involved in SCF-dependent ubiquitin-mediated proteolysis, two ubiquitin-related proteins, and a proteasome-substrate-size regulator. Vesicular trafficking is supported by four genes, including two vacuolar protein sorting-associated proteins, the µ1 subunit of the adaptor protein complex AP-2, and an intraflagellar transport component. Gene expression regulation is represented by three genes acting at the post-transcriptional level, including factors involved in RNA splicing and pre-mRNA branch point binding, and an adapter mediating XPO1/CRM1-dependent nuclear export of the 60S ribosomal subunit. Another three proteins belong to the mitochondrial respiratory chain, namely a mitochondrial carrier protein and two subunits of mitochondrial complex I.

The remaining annotated species-specific proteins do not cluster into any coherent functional category. These include a K02A2.6-like protein, a Raptor-like component of the mTOR pathway, a protein with NAD(P)-binding Rossmann-like domain, a trans-membrane Fragile-X-F protein, an acetylgalactosaminyltransferase, a Fringe-like glycosyl-transferase, a cadherin-repeat-containing protein, a pyruvate kinase, as well as proteins of unknown function (DUF3808 and DUF4485).

Twenty-four gene families were significantly contracted only in *A. incompertus*, of which five are annotated with functions related to innate immunity and developmental processes, consistent with reduced individual-level defence potentially offset by colony-level social immunity (Table S4).

In contrast to findings for eusocial species in Hymenoptera and Blattodea, we found no evidence for an expansion of chemoreceptor genes in *A. incompertus* (49 ORs & 27 IRs) or in species with alloparental care (mean: 48.1 ORs & 30.4 IRs) compared to all other analysed species (mean: 66.9 ORs & 32.3 IRs).

### Increased protein evolution with increasing social complexity

Concatenated branch dN/dS (*ω*), as a measurement of mode and strength of selection, is low and uniform across the phylogenetic tree (0.030-0.172; Fig. S8), indicating strong purifying selection. We observed highest dN/dS values on terminal branches of *A. incompertus*, *P. cylindrus* and both *Ips* species, as well as on the ancestral branch of both *Euwallacea* species (Fig. S8). However, the *Ips* and *Euwallacea* ancestral values may be unreliable due to extremely short branch lengths.

To test whether *ω* systematically differs by care phenotype, we ran several branch models for each of the 1056 single copy orthologs. These models allowed *ω* to vary among different groups of branches to test alternative hypotheses. In H0, a single *ω* was inferred for all weevils and a second for outgroups. All other alternative models tested the likelihood of two *ω* values among weevil branches on which a trait existed or not. The tested traits were egg attendance (H_EA), larval attendance (H_LA), alloparental care (H_AC), and sterile workers (H_SW). After correcting for multiple testing, the alternative hypotheses differed significantly from H0 for 506 (H_EA), 642 (H_LA), 709 (H_AC), and 665 (H_SW) of the 1056 orthogroups. In the majority of these cases, *ω* was higher on branches with the tested care behaviour compared to those without that trait (94.9%, 96.7%, 97.6%, 99.4% for the traits EA, LA, APC, and SW, respectively). These tests indicate higher rates of protein evolution within species that perform care behaviour, with an apparent increasing effect along a sociality gradient from just egg attendance towards egg attendance plus larval attendance and alloparental care. We only have a single branch on which sterile workers evolved, which decreases the power of testing its additional effect on *ω*.

### Relaxation of selection is strongest in the eusocial *A. incompertus*

We investigated a potential association of increasing social complexity in weevils with a relaxation of selection, as previously reported for Hymenoptera and termites (Ewart et al. 2024; Weyna & Romiguier 2021). For this we calculated the selection intensity parameter, k, per branch with the HyPhy RELAX method (Pond et al. 2005). A log2(k) value above zero indicates intensification of selection, while a log2(k) value below zero indicates relaxation of selection. Overall, median log2(k) was relatively variable across the phylogenetic tree, with the eusocial *A. incompertus* showing the strongest signal of relaxed selection (log2(k) = -5.58), followed by the facultatively eusocial platypodine species *P. cylindrus* (log2(k) = -2.84) (Fig. S9). Accordingly, we found high proportions of single copy orthologues under significant relaxed selection both on terminal branches (*A. incompertus*: 35.6%; *P. cylindrus*: 34.6% Fig. 6) and on the ancestral Platypodinae branch (19.3%; Fig. S10), where multiple social traits evolved. Proportions of single copy orthologues under significant intensified selection are overall comparatively low with no apparent relationship to sociality level observable (Fig. 6, S10).

**Figure 6:**
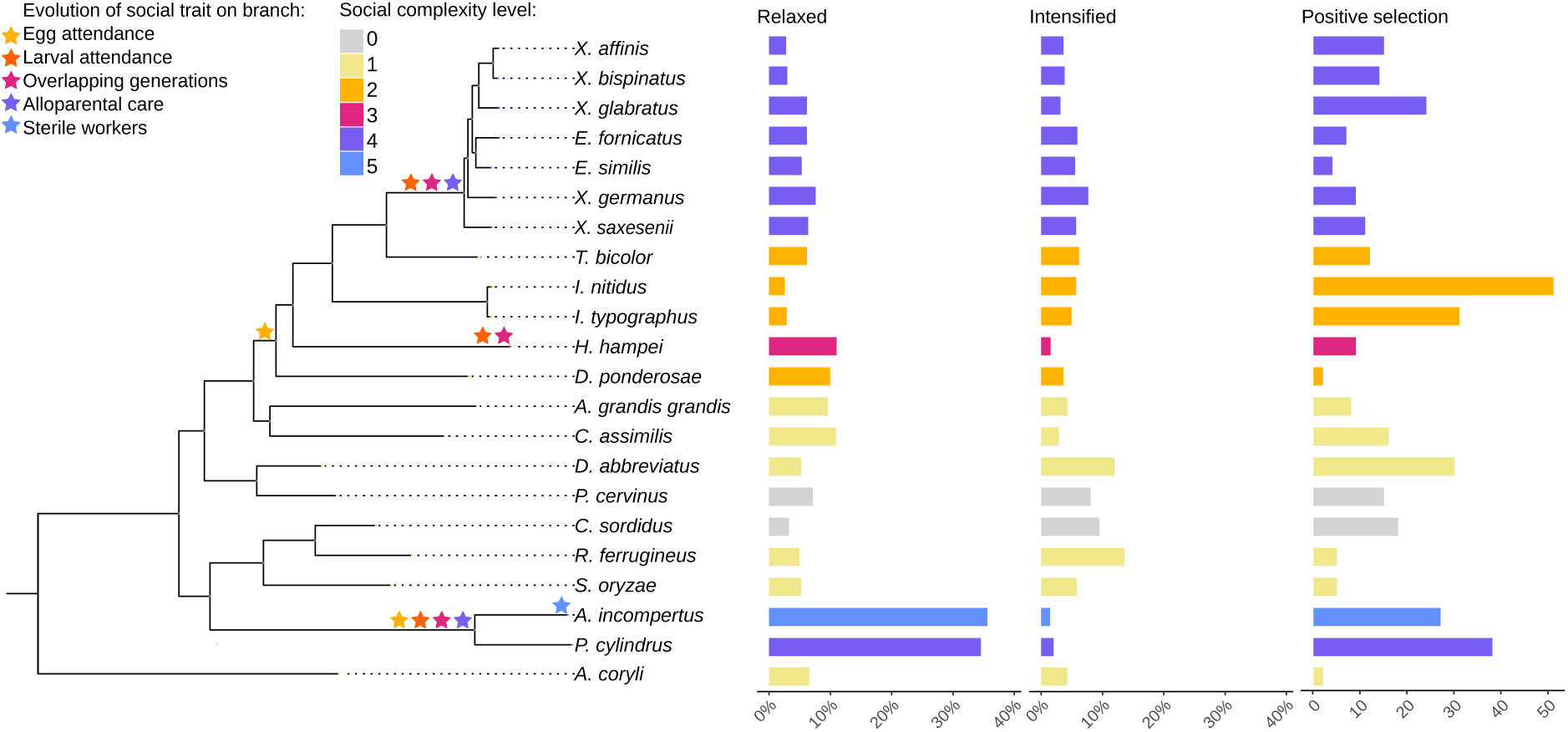
Proportion of single copy orthologues under relaxed or intensified selection for each species. Count of genes under positive selection for each species. Couloured bars indicate social complexity level: 0 = no social trait, 1 = nestbuilding only, 2 = nest building + egg attendance, 3 = nest building + egg attendance + larval attendance + overlapping generations, 4 = nest building + egg attendance + larval attendance + overlapping generations + alloparental care, and 5 = nest building + egg attendance + larval attendance + overlapping generations + alloparental care + sterile workers. Coloured stars indicate which social phenotype evolved on which branch: yellow (one star on ancestral branch) -egg attendance, orange and pink (two stars) -larval attendance and overlapping generations, orange, pink and purple (three stars) -larval attendance, overlapping generations and alloparental care, blue (one star on *A. incompertus* branch) -sterile workers.

To investigate which factors play a role in the intensification and relaxation of selection across the phylogenetic tree, we used PGLS modelling. We compared models with varying numbers of variables and two different correlation structures, namely Brownian Correlation Structure and Martins and Hansen’s (1997) covariance structure. The model with the best fit is:

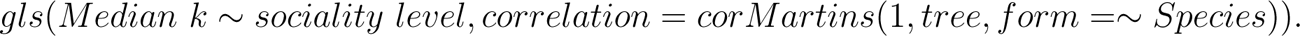

There are no PGLS models with interactions between parental care and care pattern, as the two factors have a high co-linearity. There is no significant difference between the relaxation and intensification parameter (median k) of any sociality levels, but the p-value of the comparisons between eusociality (sw) and no care (no)/nest building (nb) are close to the significance threshold of p = 0.05 (see Tables S9, S8, S7). Due to relatively few independent origins of each social trait and only one eusocial species, this model lacks power to differentiate between the effect of and interaction between different factors on median k. Nevertheless, we detected a strong standardised effect size (*>* 0.8 SD units, phylogenetically corrected) on *k* of species that perform egg attendance (social complexity level, SC: 2), alloparental care (SC: 4) or with sterile workers (SC: 5) compared to those exhibiting no care or just nest building (SC: 0 & 1; Table 3).

**Table 3:** Effect size of social complexity levels on relaxation of selection. Shown are the phylogentically corrected, standardised effect sizes (SD units) for species of the social complexity (SC) levels 2-5 compared to species that perform only nest building (SC: 1) or no care behaviours (SC: 0). Values are calculated by PGLS on standardised k values. We present also the highest social trait for each SC level -consult Fig. 1 for more details on SC levels.

| SC Level | most complex trait | nest building only | no care |
| --- | --- | --- | --- |
| 2 | egg attendance | -0.84 | -0.86 |
| 3 | overlapping generations | -0.59 | -0.61 |
| 4 | alloparental care | -1.00 | -1.02 |
| 5 | sterile workers | -2.75 | -2.78 |

### Developmental and regulatory genes under positive selection at evolutionary origins of social traits

We tested the expectation that positive selection on protein coding genes becomes more abundant with the evolution of increasing social complexity (Rehan & Toth 2015) with the HyPhy method aBSREL. We found limited support for this expectation. Although, *A. incompertus*, the most socially advanced species in our dataset, had a relatively high number of genes under significant positive selection (27), this was only fifth highest behind *I. nitidus* (51), *P. cylindrus* (38), *I. typographus* (31) and *D. abbreviatus*(30), three of which do not even perform larval attendance (Fig. S11).

At the root of Scolytinae, where egg attendance evolved, we found ten genes under positive selection, that can be functionally clustered into the functional categories: gene expression regulation, neural development, actin/cytoskeleton regulation, signaling and other functions (Table S10). On the *H. hampei* branch, where larval attendance and overlapping generations emerged, there were nine genes under positive selection with the functions oogenesis/embryogenesis, transcriptional regulation, and membrane transport. At the root of the Xyleborini clade, where larval attendance and alloparental care evolved, a single gene was under positive selection: the JNK transcription factor EGR1. At the second origin of larval attendance and alloparental care in our dataset - the Platypodinae ancestor -we identified six genes under positive selection, which can be functionally grouped as gene expression regulation, nervous system development, oogenesis/embryogenesis, and a muscle structural protein. The 27 genes under positive selection in *A. incompertus*, which may therefore be related to the evolution of sterile castes, are functionally diverse, but can be roughly grouped into the following categories: neural development, transcriptional regulation, DNA repair, membrane transport, muscle specification and cytoskeletal components as well as further, largely uncharacterised genes (Table S10).

## Discussion

In this study we sequenced and assembled three new high-quality weevil genomes, including *A. incompertus*, the only obligately eusocial beetle. With these and 18 publicly available weevil genomes we investigated the molecular mechanisms underlying the evolution of sociality in weevils. Spanning two origins of egg attendance, three origins of larval attendance and overlapping generations, two origins of alloparental care and the single origin of sterile workers within Curculionidae, this dataset offers a valuable comparative platform for the investigation of molecular mechanisms of the evolution of (eu)sociality outside of Hymenoptera and Blattodea. Hence, this platform allows for testing expectations and hypotheses generated within Hymenoptera and Blattodea, such as an increase in relaxation of selection or positive selection with increasing social complexity.

### Social weevil genomes are compact and TE-poor, with low substitution rates

As already established in most eukaryotic clades (Marino et al. 2025), we found a strong positive correlation between repetitive content and genome size within weevils. Additionally, both repetitive content and genome size tend to decrease with increasing sociality. The obligately eusocial *A. incompertus* has the smallest genome (0.1 Gb) and the lowest repetitive content (19%) in the dataset, while facultatively eusocial species in the Xyleborini and Platypodinae clades also have very small genomes (0.1-0.2 Gb), but a more variable repetitive content (22-44%). This parallels the previous finding of a negative correlation between TE content and social complexity in bees (Kapheim et al. 2015). However, general relationships between TE content and genome size and sociality appear to be lineage specific (Mikhailova et al. 2024). Interestingly, we also found protein coding content to be reduced in social weevils. *A. incompertus* has the second lowest number of protein coding genes (12034), with all facultative eusocial species having under 15000 protein coding genes, except for *X. affinis* (32381) and *X. glabratus* (41176).

With increasing social complexity two competing mechanisms may lead to an increase or decrease in evolutionary rates. Longer generation times of social insect species are expected to decrease overall substitution rates due to lower germ line replication rate per year (Lewin & Eyre-Walker 2025). Large eusocial colonies, on the other hand, also have much higher rates of reproductive output, leading to a positive correlation between colony size (a proxy for social complexity) and substitution rates (Rubin 2022). In weevils, we found a negative association between social complexity and both synonymous and non-synonymous substitution rates, which is most evident in species that perform alloparental care. Social weevils live in relatively small groups so increased generation time caused by delayed dispersal is likely causing this reduction in mutation rates, while high productivity associated with large eusocial colonies plays a lesser role.

### Evidence for sheltering and reproductive skew reducing selection efficacy

At least two traits related to sociality in insects are predicted to lead to a relaxation of purifying selection. One is the decrease in effective population size caused by reproductive skew, which leads to a decrease in selection efficacy as genetic drift becomes stronger (Ewart et al. 2024; Roux et al. 2024; Weyna & Romiguier 2021). In many ambrosia beetles, female-biased sex ratios and inbreeding can further affect effective population size and thus selection efficacy (Keller et al. 2011). The other mechanism is sheltering, where adults take care of larvae, increasing their chances of survival even if they are not well adapted (Mashoodh et al. 2023; Pascoal et al. 2023; Rinke et al. 2026). When purifying selection on individual gene family members is relaxed, loss-of-function mutations are less efficiently purged from the population, allowing gene family size to contract over evolutionary time.

We found a strong association of the social traits alloparental care and sterile workers with relaxed selection. The genome of the eusocial weevil species, *A. incompertus*, showed the strongest signature of relaxed selection and the highest proportion of genes under relaxed selection (35.61%). Although the relationship is not significant after controlling for phylogeny, a strong effect size (*>*0.8 SD units) indicates a lack of power due to two independent origins of alloparental care and one of obligate eusociality in our dataset. A medium standardised effect size (0.59 SD units) was measured for relaxation of selection with the evolution of larval dependence and overlapping generations.

Additionally, we found a significant enrichment of convergent gene family contractions on pairs of branches where social traits evolved, while convergent expansions were under-represented. Many of these convergent gene losses occurred with the evolution of larval attendance and overlapping generations, which are unlikely to lead to a marked reduction in effective population size. Higher dN/dS ratios in species with larval attendance or, especially, alloparental care offered further support for a relaxation of selection with increasing social complexity.

These results support the existence of two mechanisms leading to a relaxation of selection not only in highly social species that perform alloparental care but also in species that only carry out parental care of larval offspring. Since we expect little effect of larval attendance on reproductive skew, the medium effect of this care trait on relaxed selection, combined with multiple convergent gene losses offers strong support for the effect of sheltering, i.e. relaxed selection on traits compensated by parental care. The strongest relaxation of selection in facultatively (alloparental care) and obligately eusocial species (sterile workers) may be caused by a greater larval dependence combined with the additional effect of an increasing reproductive skew reducing the effective population size.

### Convergent gene loss as a consequence of social living

Genes under relaxed selection and contracted gene families on branches where sociality evolved share similar functions. This overlap between the two analyses supports the conclusion that these genes are correlated with the evolution of sociality. As these gene functions are lost or reduced in social species compared to non-social species, this likely reflects a consequence of social living rather than a driver of its evolution, as these functions become dispensable in a social context. Protected lifestyles inside galleries, social immunity, mutualistic fungi, and parental care with sheltering of larvae can all reduce the need for certain traits, relaxing selection on the genes underlying them. Where larval attendance and overlapping generations evolved, these functions were related to proteolysis. With the evolution of alloparental care, primary metabolism (carbohydrate and fat) and transport functions predominate. In *A. incompertus*, the only obligate eusocial weevil with sterile workers, the overlap lies in DNA metabolic processes as well as regulatory and signalling functions. Interestingly, stress response was the only function present in gene families convergently contracted on all combinations of branches where matching social phenotypes evolved. This points to a reduced need for stress response genes in larvae that receive parental protection. These results build on the findings of two previous studies in the subsocial burying beetle *Nicrophorus vespilloides* that detected increased genetic variation due to relaxed selection as an effect of parental care (Mashoodh et al. 2023; Pascoal et al. 2023). This increased genetic variation reduces offspring fitness when parental care is absent, but is alleviated when parental care is present. A reduction in fitness due to sheltering is likely to lead to increased larval dependence and a unidirectional evolution towards higher sociality.

Taken together, these findings suggest that social weevils that rear their offspring in protected galleries live in a mild environment with less extreme temperatures, predators and pathogens (due to cleaning behaviour), potentially also with reduced plant defence chemicals and less variable food sources (beetle species-specific, obligate mutualistic fungi). Consequently, genes that mediate stress responses (Hsp70), detoxification (carboxylesterases, aldo/keto-reductases, P450s), and broad carbohydrate uptake (sugar transporters) become less essential and are repeatedly lost. The repeated contraction of cysteine and serine proteases (peptidase C1, S1, M1) may be related to a shift in nutrition from digesting different plant parts themselves to symbiont-based nutrition. Ancestral Curculionidae species likley possessed a broad repertoire of stress-response, digestive and detoxifying enzymes to survive in an environment with variable, chemically defended host plants. Due to social behaviour such as parental care, gallery construction and fungal farming, a sheltered niche was constructed and genes were lost.

With these results in mind, it is important to note that sociality and gallery-dwelling fungus farming are highly collinear in this dataset. Feeding on fungus rather than wood could explain some of the observed patterns, such as the convergent losses of detoxification-related genes, meaning the effects of sociality and nutrition cannot easily be disentangled. An important exception is *H. hampei*, which shows parental care and overlapping generations but does not farm fungus and instead feeds on toxic coffee berries. Nevertheless, the gene families convergently contracted between *H. hampei* and the two other branches in which larval attendance and overlapping generations evolved shared similar functional annotations, including detoxification. However, every highly social species in this dataset, and in weevils more broadly, farms fungus. Sociality most likely evolved in a close feedback with fungus farming (Biedermann & Rohlfs 2017), since it is difficult to maintain a fungal garden alone. This raises the question of whether disentangling the effects of sociality and nutrition is even meaningful, or whether the two should instead be treated as a single, jointly evolved trait.

### Limited role of positive selection

Some studies in Hymenoptera found that positive selection plays an increasingly important role in the evolution of more complex sociality (Kapheim et al. 2015; Roux et al. 2014; Shell et al. 2021). However, in our weevil dataset, there was no clear pattern of the number of genes under positive selection across sociality levels. The low number of genes under positive selection in social species within out dataset may be due to the lack of morphological differences between reproductives and workers/carers. In weevils, workers and queens are not clearly differentiated morphologically, with workers appearing to be simply unmated individuals. Compared to the highly specialised, foraging workers within large ant colonies, for example, weevil workers remain within the gallery, mainly tending to offspring and fungus. Genes under positive selection in the obligate eusocial *A. incompertus* were broadly annotated as nervous system development, transport, and transcriptional regulation, which may play a role in the worker behaviour, rather than increased communication. Additionally, *A. incompertus* is unusual in that it inhabits living trees rather than dead wood, which may impose distinct selective pressures that contribute to the positive selection signal observed in this species. Worker phenotypes may develop through changes in genes already present (e.g. juvenile hormone signaling, nutrition-dependent development) rather than novel adaptations.

### Conclusions

In summary, our comparative genomic analysis of 21 weevil species, including the first high-quality genome of the obligate eusocial *A. incompertus*, reveals that increasing social complexity is consistently associated with smaller, less repetitive genomes, reduced substitution rates, and contractions of gene families, particularly in DNA metabolic processes, regulatory/signaling functions, primary metabolism, transport functions, and proteolytic pathways. Convergent gene family contractions were significantly enriched on branches where social traits evolved, with the strongest signatures of relaxed selection and highest dN/dS ratios in lineages with alloparental care and sterile workers, consistent with two complementary mechanisms: sheltering, whereby parental care buffers individuals against otherwise deleterious loss-of-function mutations, and a reduction in effective population size driven by reproductive skew. In contrast, neither positive selection nor extensive gene-family expansions appear to drive the evolution of weevil eusociality, contrasting with patterns reported for Hymenoptera. These results support a model in which the protected gallery environment with mutualistic fungi renders many ancestral metabolic and defensive genes dispensable, leading to convergent gene family contractions across independent origins of sociality. Expanding this framework with additional high-quality genomes and functional assays will be essential to disentangle the relative contributions of life-history, ecological niche, and demographic effects to the genomic architecture of sociality in beetles.

## Materials and Methods

### Obtaining data

We sequenced and assembled three weevil species specifically for this study, namely *A. incompertus*, *X. germanus* and *X. saxesenii*. Additionally, all available published genomes of weevil species with social phenotypes and a dataset from a previous publication (Rinke et al. 2026) were used in this analysis.

There are four batches of data:

1. Genomes of three weevil species (*A. incompertus*, *X. germanus* and *X. saxesenii*) were assembled and annotated (see Genome assembly and Genome annotation).
2. Published genomes of 13 weevil species and one leaf-rolling weevil used in Rinke et al. 2026(see Additional species, Table S1).
3. Genomes of *X. affinis*, *X. bispinatus* and *X. glabratus* were downloaded on 23.06.2025 from NCBI and re-annotated (see Genome annotation, Table S1).
4. Genomes and annotations of *E. fornicatus* and *E. similis* were downloaded from NCBI on 22.08.2025. (see Additional species, Table S1).

### Genome assembly

#### Assembly of A. incompertus

The sample used for sequencing was a single *A. incompertus* male collected in Dampier State Forest, New South Wales, Australia, in 2016 and stored at -80°C. Fresh samples were not found in 2023, likely due previous forest fires (Saunders et al. 2021). DNA was sequenced with ONT PromethION for long-read sequencing and Illumina for polishing. For the genome assembly, a frame-shift correction for long reads was performed using proovframe v.0.9.8 (Hackl et al. 2021) with diamond v.2.1.10.164 (Buchfink et al. 2021). Genome size, heterozygosity and repeat content were estimated using jellyfish v.2.3.0 (Marçais & Kingsford 2011) and GenomeScope version 1.0 (Vurture et al. 2017). Long read assembly was performed with Flye v.2.9.5 (Kolmogorov et al. 2019) and its quality checked with QUAST v.5.3.0 (Mikheenko et al. 2018) and BUSCO v.5.1.2 (Manni et al. 2021). Additional polishing of the assembly with short reads was performed using HyPo v.1.0.3 (Kundu et al. 2019) with bwa v.0.7.15 (Li 2013), minimap2 v.2.24 (Li 2021) and SAMtools v.1.16.1 (Danecek et al. 2021). Purging was performed using Purge Haplotigs v.1.1.3 (Roach et al. 2018), with cutoff values of 3, 15, and 195. Adapter and contamination check and removal was performed with fcs v.0.5.4 (Astashyn et al. 2024).

#### Assembly of X. germanus and X. saxesenii

Samples of *X. germanus* and *X. saxesenii* were collected in 2023 at the experimental forest of the chair of Forest Entomology and Protection in Stegen, Wittental, close to Freiburg, Germany, and sequenced using PacBio HiFi (Hon et al. 2020). Pre-assembly statistics were obtained using jellyfish v.2.3.0 (Marçais & Kingsford 2011). Hifiasm v.0.19.7 (Cheng et al. 2021) was used for genome assembly and compleasm v.0.2.7 (Huang & Li 2023) with arthropoda dataset, QUAST v.5.3.0 (Mikheenko et al. 2018) and Merqury v.1.3 (Rhie et al. 2020) for quality control. Adapter and contamination check and removal was performed with fcs v.0.5.4 (Astashyn et al. 2024).

### Additional species

Genomes of 13 weevil species and one leaf-rolling weevil species were re-annotated using the fasTE pipeline (Bell et al. 2022) and BRAKER2 v.3.0.8 (Brůna et al. 2021; Brůna et al. 2020; Buchfink et al. 2015; Gotoh 2008; Hoff et al. 2016; Hoff et al. 2019; Iwata & Gotoh 2012; Lomsadze et al. 2005; Stanke et al. 2008; Stanke et al. 2006) for a previous study (for more detail see Materials and Methods section of Rinke et al. 2026). The species used in that study are: *Apoderus coryli* (Crowley 2021), *P. cylindrus* (Barclay et al. 2024), *H. hampei* (Navarro-Escalante et al. 2021), *I. nitidus* (Wang et al. 2023), *I. typographus* (Powell et al. 2021), *T. bicolor* (Telfer & Badham 2024) and *D. ponderosae* (Keeling et al. 2022), *D. abbreviatus* (Sylvester et al. 2024), *R. ferrugineus* (Dias et al. 2021) and *S. oryzae* (Parisot et al. 2021), *A. grandis grandis* (Childers et al. 2021), *C. assimilis* (Pest Genomics Initiative between Rothamsted Research, Bayer, and Syngenta), *C. sordidus* (Rodriguez Ruiz & Van Dam 2023), and *P. cervinus* (Barclay et al. 2023). Additionally to these genomes used in the previous study, here, the genomes of *X. affinis*, *X. bispinatus* and *X. glabratus* (all three published by Iridian Genomes) were downloaded and re-annotated. For two more weevil species we downloaded genome and annotation files, namely *E. fornicatus* and *E. similis* (Bickerstaff et al. 2024). All genome assemblies used in this study had a very high compleasm completeness score >95% (see Table S2). Please refer to Fig. 1 for phenotypic differences between these species.

### Genome annotation

All genomes assembled in this study as well as the genomes of *X. affinis*, *X. bispinatus* and *X. glabratus* were repeat annotated using the Earlgrey v.6.3.1 (Baril et al. 2024) to softmask repeats in the genome. Then, BRAKER2 v.3.0.8 (Brůna et al. 2021; Brůna et al. 2020; Buchfink et al. 2015; Gotoh 2008; Hoff et al. 2016; Hoff et al. 2019; Iwata & Gotoh 2012; Lomsadze et al. 2005; Stanke et al. 2008; Stanke et al. 2006) was used with the arthropoda dataset to annotate protein coding genes within the genomes. The quality of the annotations was checked using compleasm v.0.2.7 (Huang & Li 2023) with arthropoda dataset. All annotations had very high completeness scores of >90% (see Table S2). The proportion of repetitive content was calculated from the soft-masked genomes, counting masked bases as repetitive content and unmasked bases as non-repetitive.

### Data preparation

To obtain proteome sequence and coding sequence (cds) fasta files, we used the gffread program from the cufflinks v.2.2.1 suite (Trapnell et al. 2010) with both, genome fasta file and gff file as input. Sequence-Fairies v.0.9.1 (Kemena 2024) seqCheck was used to remove sequences with premature stop codons and isoformCleaner was used to extract only the longest isoform of each protein. Again, compleasm v.0.2.7 (Huang & Li 2023) with arthropoda dataset was used for quality check of the proteome. To annotate protein domains, we used interproscan v.5.75 (Jones et al. 2014) with the pfam database v.106.0 (Paysan-Lafosse et al. 2025). We filtered out pfam domains associated with transposable elements, as previous analyses have shown that they interfere with the analysis of gene family size evolution. Similarly to (Rinke et al. 2026), all genes containing the following domains taken from (Min & Choi 2019) were filtered out: PF00075, PF00078, PF00665, PF02925, PF02992, PF03184, PF03221, PF03732, PF04687, PF05699, PF05840, PF05970, PF07727, PF08283, PF08284, PF10551, PF13358, PF13359, PF13456, PF13837, PF13976, PF14214, PF14223, PF14529. Number of protein coding genes was calculated once for the entire annotation and once after removing these TEs.

### Orthology and phylogenetic tree reconstruction

We ran OrthoFinder v.2.5.5 (Emms & Kelly 2019) to infer orthologous genes across all species used in this study. To prepare the data for phylogenetic tree reconstruction, all single copy orthologs were aligned using clustalo v.1.2.4 (Sievers et al. 2011) and concatenated using concatenator from the sequence-fairies v0.9.1 software suite (Kemena 2024). IQ-TREE v.3.0.1 (Wong et al. 2026) was then used for the reconstruction of the phylogenetic tree.

### Ancestral reconstruction

The evolution of social and potential confounding traits was evaluated using ancestral reconstruction. For this, we used a custom R script using a maximum likelihood approach with the libraries phytools (Revell 2024), ape (Paradis & Schliep 2019) and geiger (Pennell et al. 2014).

### Gene family size evolution

Gene family size evolution analysis is very sensitive to genes varying in copy number, for example TE-related genes. As we focus on protein coding non-TE genes, we performed an additional TE filtering step. Briefly, all orthogroups with domain annotations containing DDE, Phage integrase, PiggyBac, reverse transcriptase, YqaJ, transposase, transposition, transposable, transposon were filtered out. We used CAFE5 v.5.1.0 (Mendes et al. 2021) for the analysis of gene family size evolution across the phylogenetic tree. We first rooted the tree using the iTol Website (Letunic & Bork 2021) and made it ultrametric using the make_ultrametric.py from OrthoFinder v.2.5.4. The OrthoFinder gene count file was modified to fit the format for CAFE input files. The clade_and_size_filter.py of CAFE5 v.5.1.0 was used to filter gene families by size. First, we estimated an error model to ensure that assembly quality does not have an impact on gene family sizes. Then we tested different gamma values, *gamma* = 1 had the best likelihood score. Using the lambda from the run with *gamma* = 1, namely *lambda* = 0.218, we ran CAFE for the small and large families together. This run failed, due to the large families, so the results are based on *gamma* = 1 and not including the two families with the largest gene family size differences between the species. Also, only gene families present at the root of the tree are analysed (8833 out of 17429). CafePlotter (https://github.com/moshi4/CafePlotter) was used for plotting results. Gene families expanded and contracted on the branches of interest were extracted and overlaps between branches were plotted using a custom R script with the packages dplyr (Wickham et al. 2023) and ggvenn (Yan 2025). Gene families of interest were functionally annotated using eggnog-mapper version 2.1.12 Cantalapiedra et al. 2021; Huerta-Cepas et al. 2019 and eggNOG DB version 5.0.2, diamond version 2.1.10 Buchfink et al. 2021.

### Selection

For all selection analyses, the data were prepared by extracting the coding sequences for each single copy orthologue using sequencefairies seqExtract v.0.9.1 (Kemena 2024). Coding sequences were aligned based on the previous protein alignments using pal2nal v.14 (Suyama et al. 2006). Coding sequences were then trimmed using Gblocks v.0.91b (Castresana 2000) with parameters -t=c -b5=h. We used CODEML, part of the PAML package v.4.10.9 (Yang 2007), with a free ratios codon-substitution model (M1) to calculate dN/dS for each branch across the phylogenetic tree. For each branch, we calculated pooled dN and dS by summing N×dN and N (and separately S×dS and S) across all single copy orthologs and dividing each pooled substitution count by its pooled site count, then took the ratio of these two pooled rates to obtain branch-specific dN/dS. Branch-gene estimates with dS *≥* 3, dN/dS *≥* 10, or fewer than 10 N or S sites were excluded from the sums as unreliable. A custom R script using the packages ggtree (Xu et al. 2022; Yu 2020; Yu 2022; Yu et al. 2018; Yu et al. 2017), treeio (Wang et al. 2020), dplyr (Wickham et al. 2023), viridis (Garnier et al. 2024) and scales (Wickham et al. 2025) was used to plot the phylogenetic tree with dN/dS per branch. To test for differences in dN/dS between groups of branches, we ran multiple CODEML branch models and compared likelihood scores to the null model

To test for relaxation or intensification of selection, we used RELAX (Hyphy v.2.5.80). We ran RELAX with each branch as foreground branch, to calculate the average relaxation and intensification parameter (k) for each branch. Relax output was parsed using a custom python script and the phylogenetic tree was extracted from the output. Custom R scripts using the packages dplyr (Wickham et al. 2023), tidyverse (Wickham et al. 2019), viridis (Garnier et al. 2024), gridExtra (Auguie 2015), ape (Paradis & Schliep 2019), ggtree (Xu et al. 2022; Yu 2020; Yu 2022; Yu et al. 2018; Yu et al. 2017) and ggplot2 (Wickham 2016) were used to find orthogroups under relaxed or intensified selection and to plot median k (relaxation and intensification parameter) per branch in the phylogenetic tree, as well as density plots of k. Additionally, relax data was formatted and used in a generalized linear model as well as a PGLS using a custom R script with the packages dplyr (Wickham et al. 2023), lme4 (Bates et al. 2015), readr (Wickham et al. 2024), ggeffects (Lüdecke 2018), ggplot2 (Wickham 2016), stargazer (Hlavac 2022), emmeans (Lenth 2025), DHARMa (Hartig 2024), caret (Kuhn & Max 2008), lmerTest (Kuznetsova et al. 2017), ape (Paradis & Schliep 2019), and nlme (Pinheiro et al. 2025; Pinheiro & Bates 2000). Functions of genes under relaxed and intensified selection were annotated using the genes in *Tribolium castaneum*.

We tested for positive selection using aBSREL (Hyphy v.2.5.80) without foreground branches in an exploratory approach to calculate the number of genes under positive selection on each branch. Numbers of positively selected genes were counted for each branch using a bash script and a multiple testing correction was performed. Values were then plotted on a phylogenetic tree using a custom R script using dplyr (Wickham et al. 2023), ggplot2 (Wickham 2016), ape (Paradis & Schliep 2019), ggtree (Xu et al. 2022; Yu 2020; Yu 2022; Yu et al. 2018; Yu et al. 2017) and viridis (Garnier et al. 2024). Also the orthogoups under positive selection on the branches of interest were extracted for each foreground branch and their functions annotated using the gene annotation in *T. castaneum*.

### Substitution rates

Synonymous and non-synonymous substitution rates (dS and dN) were calculated with the FitMG94.bf script in the HyPhy suite (Pond et al. 2005) on all cds alignments with the parameters: –shared-parameters alpha,beta –frequencies CF3×4. The relationship to social traits was estimated with PGLS as with RELAX, above.

## Acknowledgements

We thank Heiko Vogel for his help with DNA extraction and sample handling.

## Data Availability

The genome assemblies produced in this study are available at NCBI (accession numbers see Table S1). Genome annotations, namely coding sequences (cds) and proteomes produced in this study are available at https://doi.org/10.5281/zenodo.22808978.

## References

1. Astashyn, A., E. S. Tvedte, D. Sweeney, V. Sapojnikov, N. Bouk, V. Joukov, E. Mozes, P. K. Strope, P. M. Sylla, L. Wagner et al. (2024). ‘Rapid and sensitive detection of genome contamination at scale with FCS-GX’. In: Genome Biol 25, p. 60. doi: 10.1186/s13059-024-03198-7.

2. Auguie, B. (2015). gridExtra: Miscellaneous Functions for “Grid” Graphics.

3. Barclay, M. V. L., Natural History Museum Genome Acquisition Lab, Darwin Tree of Life Barcoding collective, Wellcome Sanger Institute Tree of Life programme, Wellcome Sanger Institute Scientific Operations: Sequencing Operations collective, Tree of Life Core Informatics collective & Darwin Tree of Life Consortium (2023). ‘The genome sequence of a weevil, *Polydrusus cervinus* (Linnaeus, 1758)’. In: Wellcome Open Res 8, p. 563. doi: 10.12688/wellcomeopenres.20414.1.

4. Barclay, M. V. L., D. Vassiliades, W. Bayfield Farrell, J. Cristóvão, K. Matsumoto, M. Geiser, M. G. Telfer, Natural History Museum Genome Acquisition Lab, University of Oxford and Wytham Woods Genome Acquisition Lab, Darwin Tree of Life Barcoding collective et al. (2024). ‘The genome sequence of the oak pinhole borer, *Platypus cylindrus* (Fabricius, 1792)’. In: Wellcome Open Res 9, p. 305. doi: 10.12688/wellcomeopenres.22425.1.

5. Baril, T., J. Galbraith & A. Hayward (2024). ‘Earl Grey: A Fully Automated User-Friendly Transposable Element Annotation and Analysis Pipeline’. In: Mol. Biol. Evol. 41, msae068. doi: 10.1093/molbev/msae068.

6. Bates, D., M. Mächler, B. Bolker & S. Walker (2015). ‘Fitting Linear Mixed-Effects Models Using lme4’. In: J. Stat. Softw. 67, pp. 1–48. doi: 10.18637/jss.v067.i01.

7. Bell, E. A., C. L. Butler, C. Oliveira, S. Marburger, L. Yant & M. I. Taylor (2022). ‘Transposable element annotation in non-model species: The benefits of species-specific repeat libraries using semi-automated EDTA and DeepTE de novo pipelines’. In: Mol. Ecol. Resour. 22, pp. 823–833. doi: 10.1111/1755-0998.13489.

8. Bickerstaff, J. R. M., B. H. Jordal & M. Riegler (2025). ‘Biogeography and Social Family Structure Contribute to Cryptic Genomic Divergence in the Only Obligate Eusocial Beetle Species, *Austroplatypus incompertus* (Curculionidae: Platypodinae)’. In: Mol. Ecol. 34, e70076. doi: 10.1111/mec.70076.

9. Bickerstaff, J. R. M., S. S. Smith, D. S. Kent, R. A. Beaver, A. E. Seago & M. Riegler (2020). ‘A review of the distribution and host plant associations of the platypodine ambrosia beetles (Coleoptera: Curculionidae: Platypodinae) of Australia, with an electronic species identification key’. In: Zootaxa 4894, pp. 69–80. doi: 10.11646/zootaxa.4894.1.3.

10. Bickerstaff, J. R. M., T. Walsh, L. Court, G. Pandey, K. Ireland, D. Cousins, V. Caron, T. Wallenius, A. Slipinski, R. Rane et al. (2024). ‘Chromosome Structural Rearrangements in Invasive Haplodiploid Ambrosia Beetles Revealed by the Genomes of *Euwallacea fornicatus* (Eichhoff) and *Euwallacea similis* (Ferrari) (Coleoptera, Curculionidae, Scolytinae)’. In: Genome biol. evol. 16, evae226. doi: 10.1093/gbe/evae226.

11. Biedermann, P. H. W., K. D. Klepzig & M. Taborsky (2011). ‘Costs of delayed dispersal and alloparental care in the fungus-cultivating ambrosia beetle *Xyleborus affinis* Eichhoff (Scolytinae: Curculionidae)’. In: Behav. Ecol. Sociobiol. 65, pp. 1753–1761. doi: 10.1007/s00265-011-1183-5.

12. Biedermann, P. H. W. & J. A. Nuotclà (2020). ‘Social Beetles’. In: Encyclopedia of Social Insects. Ed. by C. K. Starr. Cham: Springer International Publishing, pp. 1–8. doi: 10.1007/978-3-319-90306-4_108-1.

13. Biedermann, P. H. W. & M. Rohlfs (2017). ‘Evolutionary feedbacks between insect sociality and microbial management’. In: Curr. Opin. Insect Sci. 22, pp. 92–100. doi: 10.1016/j.cois.2017.06.003.

14. Blomquist, G. J., R. Figueroa-Teran, M. Aw, M. Song, A. Gorzalski, N. L. Abbott, E. Chang & C. Tittiger (2010). ‘Pheromone production in bark beetles’. In: *Insect Biochem*. Mol. Biol. 40, pp. 699–712. doi: 10.1016/j.ibmb.2010.07.013.

15. Boomsma, J. J. & R. Gawne (2018). ‘Superorganismality and caste differentiation as points of no return: how the major evolutionary transitions were lost in translation’. In: Biol. Rev. 93, pp. 28–54. doi: 10.1111/brv.12330.

16. Brůna, T., K. J. Hoff, A. Lomsadze, M. Stanke & M. Borodovsky (2021). ‘BRAKER2: automatic eukaryotic genome annotation with GeneMark-EP+ and AUGUSTUS supported by a protein database’. In: NAR Genom Bioinform 3, lqaa108. doi: 10.1093/nargab/lqaa108.

17. Brůna, T., A. Lomsadze & M. Borodovsky (2020). ‘GeneMark-EP+: eukaryotic gene prediction with self-training in the space of genes and proteins’. In: NAR Genom Bioinform 2, lqaa026. doi: 10.1093/nargab/lqaa026.

18. Buchfink, B., K. Reuter & H.-G. Drost (2021). ‘Sensitive protein alignments at tree-of-life scale using DIAMOND’. In: Nat Methods 18, pp. 366–368. doi: 10.1038/s41592-021-01101-x.

19. Buchfink, B., C. Xie & D. H. Huson (2015). ‘Fast and sensitive protein alignment using DIAMOND’. In: Nat Methods 12, pp. 59–60. doi: 10.1038/nmeth.3176.

20. Cantalapiedra, C. P., A. Hernández-Plaza, I. Letunic, P. Bork & J. Huerta-Cepas (2021). ‘eggNOG-mapper v2: Functional Annotation, Orthology Assignments, and Domain Prediction at the Metagenomic Scale’. In: Mol. Biol. Evol. 38, pp. 5825–5829. doi: 10.1093/molbev/msab293.

21. Castresana, J. (2000). ‘Selection of conserved blocks from multiple alignments for their use in phylogenetic analysis’. In: Mol. Biol. Evol. 17, pp. 540–552. doi: 10.1093/oxfordjournals.molbev.a026334.

22. Chak, S. T. C., S. E. Harris, K. M. Hultgren, N. W. Jeffery & D. R. Rubenstein (2021). ‘Eusociality in snapping shrimps is associated with larger genomes and an accumulation of transposable elements’. In: Proc. Natl. Acad. Sci. U. S. A. 118, e2025051118. doi: 10.1073/pnas.2025051118.

23. Cheng, H., G. T. Concepcion, X. Feng, H. Zhang & H. Li (2021). ‘Haplotype-resolved de novo assembly using phased assembly graphs with hifiasm’. In: Nat. Methods 18, pp. 170–175. doi: 10.1038/s41592-020-01056-5.

24. Childers, A. K., S. M. Geib, S. B. Sim, M. F. Poelchau, B. S. Coates, T. J. Simmonds, E. D. Scully, T. P. L. Smith, C. P. Childers, R. L. Corpuz et al. (2021). ‘The USDA-ARS Ag100Pest Initiative: High-Quality Genome Assemblies for Agricultural Pest Arthropod Research’. In: Insects 12, p. 626. doi: 10.3390/insects12070626.

25. Costa, J. T. (2006). The Other Insect Societies. Harvard University Press. doi: 10.4159/9780674271616.

26. Crowley, L. M. (2021). ‘The genome sequence of the hazel leaf-roller, Apoderus coryli (Linnaeus, 1758)’. In: *Wellcome Open Res* 6, p. 315.

27. Danecek, P., J. K. Bonfield, J. Liddle, J. Marshall, V. Ohan, M. O. Pollard, A. Whitwham, T. Keane, S. A. McCarthy, R. M. Davies et al. (2021). ‘Twelve years of SAMtools and BCFtools’. In: GigaScience 10, giab008. doi: 10.1093/gigascience/giab008.

28. Dias, G. B., M. A. Altammami, H. A. F. El-Shafie, F. M. Alhoshani, M. B. Al-Fageeh, C. M. Bergman & M. M. Manee (2021). ‘Haplotype-resolved genome assembly enables gene discovery in the red palm weevil Rhynchophorus ferrugineus’. In: Sci. Rep. 11, p. 9987. doi: 10.1038/s41598-021-89091-w.

29. Eggert, A.-K. & J. K. Müller (1997). ‘Biparental care and social evolution in burying beetles: lessons from the larder’. In: The Evolution of Social Behaviour in Insects and Arachnids. Ed. by B. J. Crespi & J. C. Choe. Cambridge: Cambridge University Press, pp. 216–236. doi: 10.1017/CBO9780511721953.011.

30. Emms, D. M. & S. Kelly (2019). ‘OrthoFinder: phylogenetic orthology inference for comparative genomics’. In: Genome Biol 20, p. 238. doi: 10.1186/s13059-019-1832-y.

31. Ewart, K. M., S. Y. W. Ho, A.-A. Chowdhury, F. R. Jaya, Y. Kinjo, J. Bennett, T. Bourguignon, H. A. Rose & N. Lo (2024). ‘Pervasive relaxed selection in termite genomes’. In: Proc Biol Sci 291, p. 20232439. doi: 10.1098/rspb.2023.2439.

32. Garnier, Simon, Ross, Noam, Rudis, Robert, Camargo, A. Pedro, Sciaini, Marco, et al. (2024). viridis(Lite) -Colorblind-Friendly Color Maps for R. doi: 10.5281/zenodo.4679423.

33. Gotoh, O. (2008). ‘A space-efficient and accurate method for mapping and aligning cDNA sequences onto genomic sequence’. In: Nucleic Acids Res 36, pp. 2630–2638. doi: 10.1093/nar/gkn105.

34. Hackl, T., F. Trigodet, A. M. Eren, S. J. Biller, J. M. Eppley, E. Luo, A. Burger, E. F. DeLong & M. G. Fischer (2021). proovframe: frameshift-correction for long-read (meta)genomics. doi: 10.1101/2021.08.23.457338.

35. Harris, J. A., K. G. Campbell & G. Wright (1976). ‘Ecological studies on the horizontal borer ’*Austroplatypus incompertus*’ (Schedl) (Coleoptera: Platypodidae)’. In: J. Aust. Entomol. Soc. 9, pp. 11–21. doi: 10.3316/informit.214637326338937.

36. Hartig, F. (2024). DHARMa: Residual Diagnostics for Hierarchical (Multi-Level / Mixed) Regression Models.

37. Hlavac, M. (2022). stargazer: Well-Formatted Regression and Summary Statistics Tables. Social Policy Institute. Bratislava, Slovakia.

38. Hoff, K. J., S. Lange, A. Lomsadze, M. Borodovsky & M. Stanke (2016). ‘BRAKER1: Unsupervised RNA-Seq-Based Genome Annotation with GeneMark-ET and AUGUSTUS’. In: Bioinformatics 32, pp. 767–769. doi: 10.1093/bioinformatics/btv661.

39. Hoff, K. J., A. Lomsadze, M. Borodovsky & M. Stanke (2019). ‘Whole-Genome Annotation with BRAKER’. In: Methods Mol Biol 1962, pp. 65–95. doi: 10.1007/978-1-4939-9173-0_5.

40. Hon, T., K. Mars, G. Young, Y.-C. Tsai, J. W. Karalius, J. M. Landolin, N. Maurer, D. Kudrna, M. A. Hardigan, C. C. Steiner et al. (2020). ‘Highly accurate long-read HiFi sequencing data for five complex genomes’. In: *Sci*. Data 7, p. 399. doi: 10.1038/s41597-020-00743-4.

41. Huang, N. & H. Li (2023). ‘compleasm: a faster and more accurate reimplementation of BUSCO’. In: Bioinformatics 39, btad595. doi: 10.1093/bioinformatics/btad595.

42. Huerta-Cepas, J., D. Szklarczyk, D. Heller, A. Hernández-Plaza, S. K. Forslund, H. Cook, D. R. Mende, I. Letunic, T. Rattei, L. J. Jensen et al. (2019). ‘eggNOG 5.0: a hierarchical, functionally and phylogenetically annotated orthology resource based on 5090 organisms and 2502 viruses’. In: Nucleic Acids Res 47, pp. D309–D314. doi: 10.1093/nar/gky1085.

43. Hulcr, J. & L. L. Stelinski (2017). ‘The Ambrosia Symbiosis: From Evolutionary Ecology to Practical Management’. In: Annu. Rev. Entomol 62, pp. 285–303. doi: 10.1146/annurev-ento-031616-035105.

44. Hunt, B. G., L. Ometto, Y. Wurm, D. Shoemaker, S. V. Yi, L. Keller & M. A. D. Goodisman (2011). ‘Relaxed selection is a precursor to the evolution of phenotypic plasticity’. In: Proc. Natl. Acad. Sci. 108, pp. 15936–15941. doi: 10.1073/pnas.1104825108.

45. Iwata, H. & O. Gotoh (2012). ‘Benchmarking spliced alignment programs including Spaln2, an extended version of Spaln that incorporates additional species-specific features’. In: Nucleic Acids Res 40, e161. doi: 10.1093/nar/gks708.

46. Jones, A. R. C., A. A. Mikhailova, C. Aumont, J. Berger, C. Liu, S. He, Z. Wang, S. Winkler, E. Bornberg-Bauer, F. Legendre et al. (2026). ‘*Cryptocercus Genomes* Expand Knowledge of Adaptations to Xylophagy and Termite Sociality’. In: Genome biol. evol. 18, evag028. doi: 10.1093/gbe/evag028.

47. Jones, P., D. Binns, H.-Y. Chang, M. Fraser, W. Li, C. McAnulla, H. McWilliam, J. Maslen, A. Mitchell, G. Nuka et al. (2014). ‘InterProScan 5: genome-scale protein function classification’. In: Bioinformatics 30, pp. 1236–1240. doi: 10.1093/bioinformatics/btu031.

48. Kapheim, K. M., H. Pan, C. Li, S. L. Salzberg, D. Puiu, T. Magoc, H. M. Robertson, M. E. Hudson, A. Venkat, B. J. Fischman et al. (2015). ‘Genomic signatures of evolutionary transitions from solitary to group living’. In: Science 348, pp. 1139–1143.

49. Keeling, C. I., E. O. Campbell, P. D. Batista, V. A. Shegelski, S. A. L. Trevoy, D. P. W. Huber, J. K. Janes & F. A. H. Sperling (2022). ‘Chromosome-level genome assembly reveals genomic architecture of northern range expansion in the mountain pine beetle, *Dendroctonus ponderosae* Hopkins (Coleoptera: Curculionidae)’. In: Mol. Ecol. Resour. 22, pp. 1149–1167. doi: 10.1111/1755-0998.13528.

50. Keller, L., K. Peer, C. Bernasconi, M. Taborsky & D. M. Shuker (2011). ‘Inbreeding and selection on sex ratio in the bark beetle Xylosandrus germanus’. In: BMC Evolutionary Biology 11, p. 359. doi: 10.1186/1471-2148-11-359.

51. Kemena, C. (2024). Sequence-Fairies. doi: 10.5281/zenodo.14288256.

52. Kent, D. S. (2008). ‘Distribution and host plant records of ’*Austroplatypus incompertus*’ (Scheld) (Coleoptera: Curculionidae: Platypodinae)’. In: Aust. J. Entomol. 35, pp. 1–6.

53. Kent, D. S. & J. A. Simpson (1992). ‘Eusociality in the beetle *Austroplatypus incompertus* (Coleoptera: Curculionidae)’. In: Naturwissenschaften 79, pp. 86–87. doi: 10.1007/BF01131810.

54. Kirkendall, L., P. H. W. Biedermann & B. Jordal (2015). ‘Evolution and Diversity of Bark and Ambrosia Beetles’. In: Bark Beetles: Biology and Ecology of Native and Invasive Species, pp. 85–156. doi: 10.1016/B978-0-12-417156-5.00003-4.

55. Kölliker, M. (2012). The Evolution of Parental Care. Ed. by N. J. Royle & P. T. Smiseth. Oxford University Press. doi: 10.1093/acprof:oso/9780199692576.001.0001.

56. Kolmogorov, M., J. Yuan, Y. Lin & P. A. Pevzner (2019). ‘Assembly of long, error-prone reads using repeat graphs’. In: Nat. Biotechnol. 37, pp. 540–546. doi: 10.1038/s41587-019-0072-8.

57. Kuhn & Max (2008). ‘Building Predictive Models in R Using the caret Package’. In: J. Stat. Softw. 28, pp. 1–26. doi: 10.18637/jss.v028.i05.

58. Kundu, R., J. Casey & W.-K. Sung (2019). HyPo: Super Fast & Accurate Polisher for Long Read Genome Assemblies. doi: 10.1101/2019.12.19.882506.

59. Kuznetsova, A., P. B. Brockhoff & R. H. B. Christensen (2017). ‘lmerTest Package: Tests in Linear Mixed Effects Models’. In: J. Stat. Softw. 82, pp. 1–26. doi: 10.18637/jss.v082.i13.

60. Lenth, R. V. (2025). emmeans: Estimated Marginal Means, aka Least-Squares Means.

61. Letunic, I. & P. Bork (2021). ‘Interactive Tree Of Life (iTOL) v5: an online tool for phylogenetic tree display and annotation’. In: Nucleic Acids Res 49, W293–W296. doi: 10.1093/nar/gkab301.

62. Lewin, L. & A. Eyre-Walker (2025). ‘Estimates of the mutation rate per year can explain why the molecular clock depends on generation time’. In: Mol. Biol. Evol. 42, msaf069.

63. Li, H. (2013). Aligning sequence reads, clone sequences and assembly contigs with BWAMEM. doi: 10.48550/arXiv.1303.3997.

64. Li, H. (2021). ‘New strategies to improve minimap2 alignment accuracy’. In: Bioinformatics 37, pp. 4572–4574. doi: 10.1093/bioinformatics/btab705.

65. Lomsadze, A., V. Ter-Hovhannisyan, Y. O. Chernoff & M. Borodovsky (2005). ‘Gene identification in novel eukaryotic genomes by self-training algorithm’. In: Nucleic Acids Res 33, pp. 6494–6506. doi: 10.1093/nar/gki937.

66. Lüdecke, D. (2018). ‘ggeffects: Tidy Data Frames of Marginal Effects from Regression Models.’ In: J. Open Source Softw. 3, p. 772. doi: 10.21105/joss.00772.

67. Manni, M., M. R. Berkeley, M. Seppey & E. M. Zdobnov (2021). ‘BUSCO: Assessing Genomic Data Quality and Beyond’. In: Curr Protoc 1, e323. doi: 10.1002/cpz1.323.

68. Marçais, G. & C. Kingsford (2011). ‘A fast, lock-free approach for efficient parallel counting of occurrences of k-mers’. In: Bioinformatics 27, pp. 764–770. doi: 10.1093/bioinformatics/btr011.

69. Marino, A., G. Debaecker, A.-S. Fiston-Lavier, A. Haudry & B. Nabholz (2025). ‘Effective population size does not explain long-term variation in genome size and transposable element content in animals’. In: *Elife* 13, RP100574.

70. Mashoodh, R., A. T. Trowsdale, A. Manica & R. M. Kilner (2023). ‘Parental care shapes the evolution of molecular genetic variation’. In: Evol Lett 7, pp. 379–388. doi: 10.1093/evlett/qrad039.

71. Mendes, F. K., D. Vanderpool, B. Fulton & M. W. Hahn (2021). ‘CAFE 5 models variation in evolutionary rates among gene families’. In: Bioinformatics 36, pp. 5516–5518. doi: 10.1093/bioinformatics/btaa1022.

72. Mikhailova, A. A., S. Rinke & M. C. Harrison (2024). ‘Genomic signatures of eusocial evolution in insects’. In: Curr. Opin. Insect Sci. 61, p. 101136. doi: 10.1016/j.cois.2023.101136.

73. Mikheenko, A., A. Prjibelski, V. Saveliev, D. Antipov & A. Gurevich (2018). ‘Versatile genome assembly evaluation with QUAST-LG’. In: *Bioinformatics (Oxford*, England) 34, pp. i142–i150. doi: 10.1093/bioinformatics/bty266.

74. Min, B. & I.-G. Choi (2019). ‘Practical Guide for Fungal Gene Prediction from Genome Assembly and RNA-Seq Reads by FunGAP’. In: Methods Mol Biol 1962, pp. 53–64. doi: 10.1007/978-1-4939-9173-0_4.

75. Mueller, R. (2019). ‘Characterisation of fungal symbionts and microbial communities of Austroplatypus incompertus (Platypodinae) and other Australian ambrosia beetle species’. PhD thesis. Penrith, N.S.W: Western Sydney University.

76. Navarro-Escalante, L., E. M. Hernandez-Hernandez, J. Nuñez, F. E. Acevedo, A. Berrio, L. M. Constantino, B. E. Padilla-Hurtado, D. Molina, C. Gongora, R. Acuña et al. (2021). ‘A coffee berry borer (*Hypothenemus hampei*) genome assembly reveals a reduced chemosensory receptor gene repertoire and male-specific genome sequences’. In: Sci. Rep. 11, p. 4900. doi: 10.1038/s41598-021-84068-1.

77. Paradis, E. & K. Schliep (2019). ‘ape 5.0: an environment for modern phylogenetics and evolutionary analyses in R’. In: Bioinformatics 35, pp. 526–528. doi: 10.1093/bioinformatics/bty633.

78. Parisot, N., C. Vargas-Chávez, C. Goubert, P. Baa-Puyoulet, S. Balmand, L. Beranger, C. Blanc, A. Bonnamour, M. Boulesteix, N. Burlet et al. (2021). ‘The transposable element-rich genome of the cereal pest *Sitophilus oryzae*’. In: BMC Biology 19, p. 241. doi: 10.1186/s12915-021-01158-2.

79. Pascoal, S., H. Shimadzu, R. Mashoodh & R. M. Kilner (2023). ‘Parental care results in a greater mutation load, for which it is also a phenotypic antidote’. In: Proc Biol Sci 290, p. 20230115. doi: 10.1098/rspb.2023.0115.

80. Paysan-Lafosse, T., A. Andreeva, M. Blum, S. R. Chuguransky, T. Grego, B. L. Pinto, G. A. Salazar, M. L. Bileschi, F. Llinares-López, L. Meng-Papaxanthos et al. (2025). ‘The Pfam protein families database: embracing AI/ML’. In: Nucleic Acids Res 53, pp. D523–D534. doi: 10.1093/nar/gkae997.

81. Pennell, M. W., J. M. Eastman, G. J. Slater, J. W. Brown, J. C. Uyeda, R. G. FitzJohn, M. E. Alfaro & L. J. Harmon (2014). ‘geiger v2.0: an expanded suite of methods for fitting macroevolutionary models to phylogenetic trees’. In: Bioinformatics 30, pp. 2216–2218. doi: 10.1093/bioinformatics/btu181.

82. Pinheiro, J., D. Bates & R Core Team (2025). *nlme: Linear and Nonlinear Mixed Effects Models*.

83. Pinheiro, J. C. & D. M. Bates (2000). Mixed-Effects Models in S and S-PLUS. New York: Springer. doi: 10.1007/b98882.

84. Pond, S. L. K., S. D. W. Frost & S. V. Muse (2005). ‘HyPhy: hypothesis testing using phylogenies’. In: Bioinformatics 21, pp. 676–679. doi: 10.1093/bioinformatics/bti079.

85. Powell, D., E. Groe-Wilde, P. Krokene, A. Roy, A. Chakraborty, C. Löfstedt, H. Vogel, M. N. Andersson & F. Schlyter (2021). ‘A highly-contiguous genome assembly of the Eurasian spruce bark beetle, *Ips typographus*, provides insight into a major forest pest’. In: *Commun*. Biol. 4, pp. 1–9. doi: 10.1038/s42003-021-02602-3.

86. Rehan, S. M. & A. L. Toth (2015). ‘Climbing the social ladder: the molecular evolution of sociality’. In: Trends Ecol Evol 30, pp. 426–433. doi: 10.1016/j.tree.2015.05.004.

87. Revell, L. J. (2024). ‘phytools 2.0: an updated R ecosystem for phylogenetic comparative methods (and other things).’ In: PeerJ 12, e16505. doi: 10.7717/peerj.16505.

88. Rhie, A., B. P. Walenz, S. Koren & A. M. Phillippy (2020). ‘Merqury: reference-free quality, completeness, and phasing assessment for genome assemblies’. In: Genome Biol 21, p. 245. doi: 10.1186/s13059-020-02134-9.

89. Rinke, S., P. H. W. Biedermann, M. Schebeck & M. C. Harrison (2026). ‘Genomic Insights Into the Evolution of Parental Care in Weevils’. In: Genome biol. evol. 18, evag142. doi: 10.1093/gbe/evag142.

90. Roach, M. J., S. A. Schmidt & A. R. Borneman (2018). ‘Purge Haplotigs: allelic contig reassignment for third-gen diploid genome assemblies’. In: BMC Bioinformatics 19, p. 460. doi: 10.1186/s12859-018-2485-7.

91. Rodriguez Ruiz, A. & A. R. Van Dam (2023). ‘Metagenomic binning of PacBio HiFi data prior to assembly reveals a complete genome of *Cosmopolites sordidus* (Germar) (Coleopterea: Curculionidae, Dryophthorinae) the most damaging arthropod pest of bananas and plantains’. In: PeerJ 11, e16276. doi: 10.7717/peerj.16276.

92. Roux, C., A. Ha, A. Weyna, M. Lode & J. Romiguier (2024). ‘The impact of social complexity on the efficacy of natural selection in termites’. In: *Peer community j*. 4.

93. Roux, J., E. Privman, S. Moretti, J. T. Daub, M. Robinson-Rechavi & L. Keller (2014). ‘Patterns of Positive Selection in Seven Ant Genomes’. In: Mol. Biol. Evol. 31, pp. 1661–1685. doi: 10.1093/molbev/msu141.

94. Rubin, B. E. R. (2022). ‘Social insect colony size is correlated with rates of molecular evolution’. In: Insectes Sociaux 69, pp. 147–157. doi: 10.1007/s00040-022-00859-3.

95. Saunders, M. E., P. S. Barton, J. R. M. Bickerstaff, L. Frost, T. Latty, B. D. Lessard, E. C. Lowe, J. Rodriguez, T. E. White & K. D. L. Umbers (2021). ‘Limited understanding of bushfire impacts on Australian invertebrates’. In: Insect Conserv. Divers. 14, pp. 285–293. doi: 10.1111/icad.12493.

96. Shell, W. A., M. A. Steffen, H. K. Pare, A. S. Seetharam, A. J. Severin, A. L. Toth & S. M. Rehan (2021). ‘Sociality sculpts similar patterns of molecular evolution in two independently evolved lineages of eusocial bees’. In: Commun. Biol. 4, p. 253.

97. Sievers, F., A. Wilm, D. Dineen, T. J. Gibson, K. Karplus, W. Li, R. Lopez, H. McWilliam, M. Remmert, J. Söding et al. (2011). ‘Fast, scalable generation of high-quality protein multiple sequence alignments using Clustal Omega’. In: *Mol*. Syst. Biol. 7, p. 539. doi: 10.1038/msb.2011.75.

98. Six, D. L. (2012). ‘Ecological and Evolutionary Determinants of Bark Beetle —Fungus Symbioses’. In: Insects 3, pp. 339–366. doi: 10.3390/insects3010339.

99. Smith, S. M. (2013). ‘Spatial and Temporal Patterns of Kin and Population Genetic Structure of the Eusocial Beetle, Austroplatypus incompertus’. PhD thesis. Macquarie University.

100. Smith, S. S., A. Beattie, D. S. Kent & A. Stow (2009). ‘Ploidy of the eusocial beetle *Austroplatypus incompertus* (Schedl) (Coleoptera, Curculionidae) and implications for the evolution of eusociality’. In: Insectes Soc. 56, pp. 285–288. doi: 10.1007/s00040-009-0022-4.

101. Smith, S. S., D. S. Kent, J. Boomsma & A. Stow (2018). ‘Monogamous sperm storage and permanent worker sterility in a long-lived ambrosia beetle’. In: *Nat*. Ecol. Evol. 2. doi: 10.1038/s41559-018-0533-3.

102. Stanke, M., M. Diekhans, R. Baertsch & D. Haussler (2008). ‘Using native and syntenically mapped cDNA alignments to improve de novo gene finding’. In: Bioinformatics 24, pp. 637–644. doi: 10.1093/bioinformatics/btn013.

103. Stanke, M., O. Schöffmann, B. Morgenstern & S. Waack (2006). ‘Gene prediction in eukaryotes with a generalized hidden Markov model that uses hints from external sources’. In: BMC bioinformatics 7, p. 62. doi: 10.1186/1471-2105-7-62.

104. Suyama, M., D. Torrents & P. Bork (2006). ‘PAL2NAL: robust conversion of protein sequence alignments into the corresponding codon alignments’. In: Nucleic Acids Res 34, W609–612. doi: 10.1093/nar/gkl315.

105. Sylvester, T., R. Adams, W. B. Hunter, X. Li, B. Rivera-Marchand, R. Shen, N. R. Shin & D. D. McKenna (2024). ‘The genome of the invasive and broadly polyphagous *Diaprepes* root weevil, *Diaprepes abbreviatus* (Coleoptera), reveals an arsenal of putative polysaccharide-degrading enzymes’. In: J Hered 115, pp. 94–102. doi: 10.1093/jhered/esad064.

106. Telfer, M. G. & X. R. Badham (2024). ‘The genome sequence of the beech bark beetle, *Taphrorychus bicolor* (Herbst, 1793)’. In: Wellcome Open Res 9, p. 213. doi: 10.12688/wellcomeopenres.21265.1.

107. Trapnell, C., B. A. Williams, G. Pertea, A. Mortazavi, G. Kwan, M. J. van Baren, S. L. Salzberg, B. J. Wold & L. Pachter (2010). ‘Transcript assembly and quantification by RNA-Seq reveals unannotated transcripts and isoform switching during cell differentiation’. In: Nat Biotechnol 28, pp. 511–515. doi: 10.1038/nbt.1621.

108. Vurture, G. W., F. J. Sedlazeck, M. Nattestad, C. J. Underwood, H. Fang, J. Gurtowski & M. C. Schatz (2017). ‘GenomeScope: fast reference-free genome profiling from short reads’. In: Bioinformatics 33, pp. 2202–2204. doi: 10.1093/bioinformatics/btx153.

109. Wang, L.-G., Tommy Tsan-Yuk Lam, S. Xu, Z. Dai, L. Zhou, T. Feng, P. Guo, Casey W. Dunn, Bradley R. Jones, Tyler Bradley et al. (2020). ‘treeio: an R package for phylogenetic tree input and output with richly annotated and associated data.’ In: Mol. Biol. Evol. 37 (2), pp. 599–603. doi: 10.1093/molbev/msz240.

110. Wang, Z., Y. Liu, H. Wang, A. Roy, H. Liu, F. Han, X. Zhang & Q. Lu (2023). ‘Genome and transcriptome of *Ips nitidus* provide insights into high-altitude hypoxia adaptation and symbiosis’. In: iScience 26, p. 107793. doi: 10.1016/j.isci.2023.107793.

111. Weyna, A. & J. Romiguier (2021). ‘Relaxation of purifying selection suggests low effective population size in eusocial Hymenoptera and solitary pollinating bees’. In: Peer community j. 1.

112. Wickham, H. (2016). ggplot2: Elegant Graphics for Data Analysis. Springer-Verlag New York.

113. Wickham, H., M. Averick, J. Bryan, W. Chang, L. D. McGowan, R. François, G. Grolemund, A. Hayes, L. Henry, J. Hester et al. (2019). ‘Welcome to the tidyverse’. In: J. Open Source Softw. 4, p. 1686. doi: 10.21105/joss.01686.

114. Wickham, H., R. François, L. Henry, K. Müller & D. Vaughan (2023). dplyr: A Grammar of Data Manipulation.

115. Wickham, H., J. Hester & J. Bryan (2024). readr: Read Rectangular Text Data.

116. Wickham, H., T. L. Pedersen & D. Seidel (2025). scales: Scale Functions for Visualization.

117. Wilson, E. O. et al. (1971). The insect societies. Cambridge, Massachusetts, USA, Harvard University Press.

118. Wong, T. K. F., N. Ly-Trong, H. Ren, P. Demotte, H. Baños, A. J. Roger, E. Susko, C. Bielow, N. De Maio, N. Goldman et al. (2026). ‘IQ-TREE 3: phylogenomic inference software using complex evolutionary models’. In: Mol. Biol. Evol. 43, msag117. doi: 10.1093/molbev/msag117.

119. Xu, S., L. Li, X. Luo, M. Chen, W. Tang, L. Zhan, Z. Dai, Tommy T. Lam, Y. Guan & G. Yu (2022). ‘Ggtree: A serialized data object for visualization of a phylogenetic tree and annotation data’. In: iMeta 1, e56. doi: 10.1002/imt2.56.

120. Yan, L. (2025). ggvenn: Draw Venn Diagram by ’ggplot2’.

121. Yang, Z. (2007). ‘PAML 4: Phylogenetic Analysis by Maximum Likelihood’. In: Mol. Biol. Evol. 24, pp. 1586–1591. doi: 10.1093/molbev/msm088.

122. Yu, G. (2020). ‘Using ggtree to Visualize Data on Tree-Like Structures’. In: Curr Protoc Bioinformatics 69, e96. doi: 10.1002/cpbi.96.

123. Yu, G. (2022). Data Integration, Manipulation and Visualization of Phylogenetic Trees. 1st edition. Chapman and Hall/CRC. doi: 10.1201/9781003279242.

124. Yu, G., T. T.-Y. Lam, H. Zhu & Y. Guan (2018). ‘Two methods for mapping and visualizing associated data on phylogeny using ggtree.’ In: Mol. Biol. Evol. 35 (2), pp. 3041–3043. doi: 10.1093/molbev/msy194.

125. Yu, G., D. Smith, H. Zhu, Y. Guan & T. T.-Y. Lam (2017). ‘ggtree: an R package for visualization and annotation of phylogenetic trees with their covariates and other associated data.’ In: Methods ecol. evol. 8 (1), pp. 28–36. doi: 10.1111/2041-210X.12628.

