## Supplementary_material for "Sociality in weevils is shaped by sheltering and convergent gene losses"

Table S1: NCBI accession numbers for each published genome. *not published yet, accession number not yet available. Bioprojects: PRJNA1328948 (*X. germanus*), PRJNA1328956 (*A. incompertus*), PRJNA1328639 (*X. saxesenii*).

| **Organism Name** | **NCBI Accession** |
|---|---|
| *Apoderus coryli* | GCA_911728435.2 |
| *Anthonomus grandis grandis* | GCF_022605725.1 |
| *Austroplatypus incompertus* | * |
| *Ceutorhynchus assimilis* | GCA_917834065.1 |
| *Cosmopolites sordidus* | GCA_031761425.1 |
| *Diaprepes abbreviatus* | GCA_034092305.1 |
| *Dendroctonus ponderosae* | GCF_020466585.1 |
| *Euwallacea fornicatus* | GCA_040115645.1 |
| *Euwallacea similis* | GCA_039881205.1 |
| *Hypothenemus hampei* | GCA_013372445.1 |
| *Ips nitidus* | GCA_018691245.2 |
| *Ips typographus* | GCA_016097725.1 |
| *Polydrusus cervinus* | GCA_935413205.1 |
| *Platypus cylindrus* | GCA_949748235.1 |
| *Rhynchophorus ferrugineus* | GCA_014462685.1 |
| *Sitophilus oryzae* | GCF_002938485.1 |
| *Taphrorychus bicolor* | GCA_951812265.1 |
| *Tribolium castaneum* | GCA_031307605.1 |
| *Xyleborus affinis* | GCA_046766355.1 |
| *Xyleborus bispinatus* | GCA_049176215.1 |
| *Xylosandrus germanus* | * |
| *Xyleborus glabratus* | GCA_049176235.1 |
| *Xyleborinus saxesenii* | * |

### Supplements

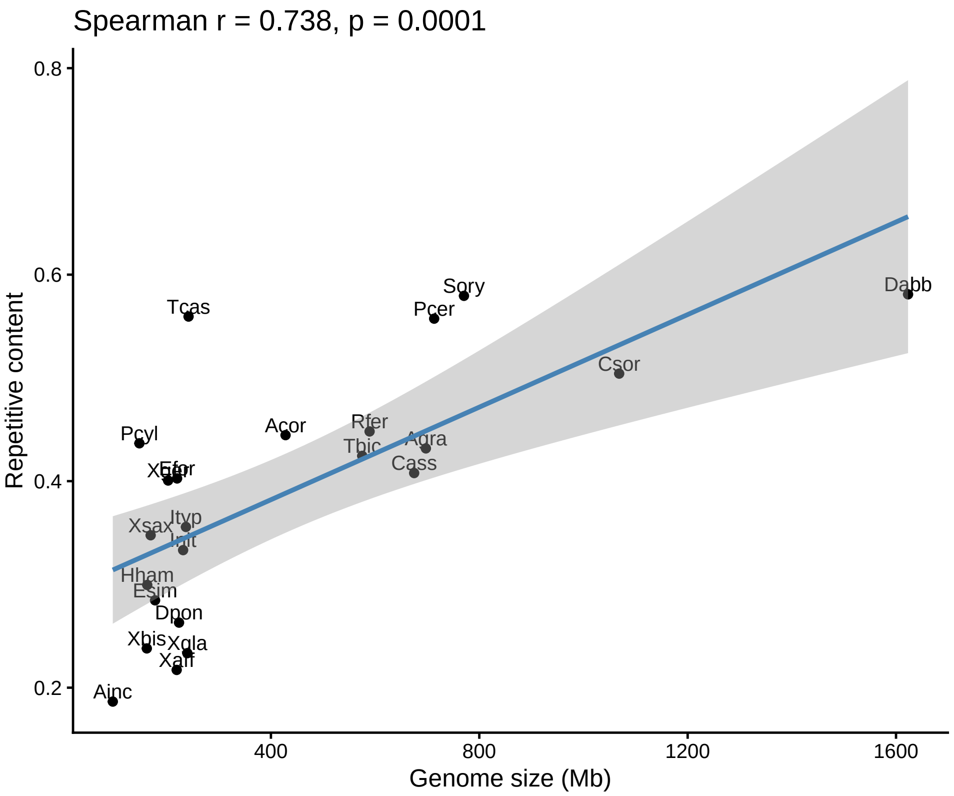


Figure S1: Repetitive content (%) was plotted against genome size (Mb). Spearman correlation was calculated and shows a strong positive correlation. Species abbreviations: Acor = *A. coryli*, Agra = *A. grandis grandis*, Ainc = *A. incompertus*, Cass = *C. assimilis*, Csor = *C. sordidus*, Dabb = *D. abbreviatus*, Dpon = *D. ponderosae*, Efor = *E. fornicatus*, Esim = *E. similis*, Hham = *H. hampei*, Init = *I. nitidus*, Ityp = *I. typographus*, Pcer = *P. cervinus*, Pcyl = *P. cylindrus*, Rfer = *R. ferrugineus*, Sory = *S. oryzae*, Tbic = *T. bicolor*, Tcas = *T. castaneum*, Xaff = *X. affinis*, Xbis = *X. bispinatus*, Xger = *X. germanus*, Xgla = *X. glabratus*, Xsax = *X. saxesenii.*

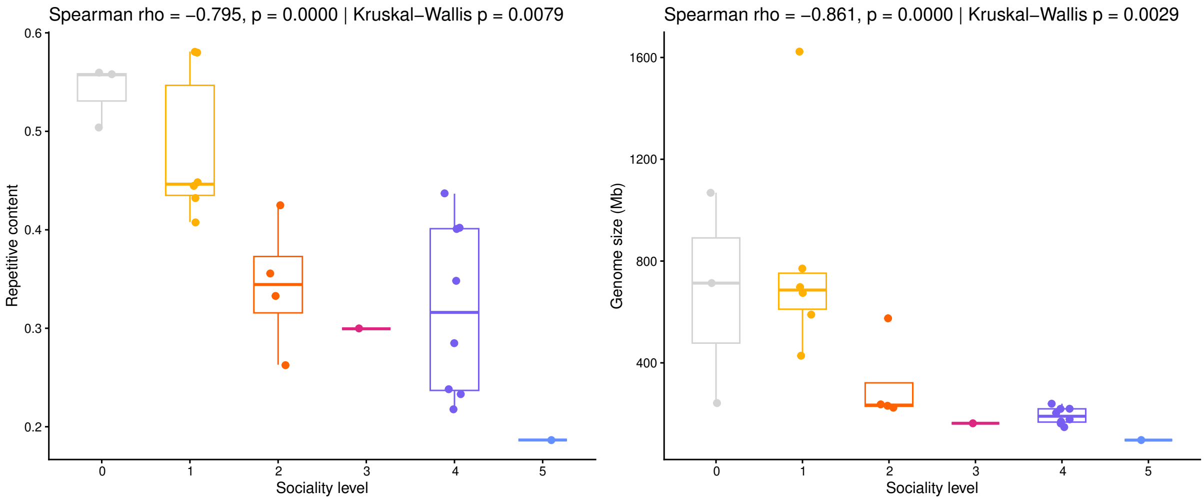


(a) Repetitive content

(b) Genome size

Figure S2: Correlations between sociality levels and a) repetitive genome content and b) genome size. 0 = no social trait, 1 = nest building only, 2 = nest building + egg attendance, 3 = nest building + egg attendance + larval attendance + overlapping generations, 4 = nest building + egg attendance + larval attendance + overlapping generations + alloparental care, and 5 = nest building + egg attendance + larval attendance + overlapping generations + alloparental care + sterile workers.

Table S2: Assembly and annotation statistics. S = complete and single copy genes (%), D = complete and duplicated genes (%), C = completeness score (%), Contigs = number of contigs, GC = GC content (%).

| | Assembly | | | | | | | | | | Annotation | | |
|---|---|---|---|---|---|---|---|---|---|---|---|---|---|
| Species | S | D | C | Contigs | Total length | GC | N50 | N90 | L50 | L90 | S | D | C |
| *A. coryli* | 97.78 | 1.80 | 99.58 | 129 | 428,077,388 | 32.67 | 23,694,900 | 12,555,284 | 6 | 16 | 89.32 | 7.26 | 96.58 |
| *A. grandis grandis* | 97.66 | 1.68 | 99.34 | 304 | 697,452,952 | 32.13 | 36,454,210 | 20,361,090 | 8 | 18 | 84.88 | 6.06 | 90.94 |
| *A. incompertus* | 95.08 | 1.38 | 96.46 | 1451 | 96,366,439 | 28.11 | 194,133 | 44,826 | 140 | 512 | 89.74 | 6.72 | 96.46 |
| *C. assimilis* | 95.38 | 2.82 | 98.20 | 44 | 674,950,870 | 35.29 | 55,466,945 | 22,725,061 | 5 | 12 | 83.50 | 8.94 | 92.44 |
| *C. sordidus* | 97.00 | 0.84 | 97.84 | 2,977 | 1,068,542,587 | 33.97 | 706,383 | 179,964 | 443 | 1557 | 89.38 | 5.10 | 94.48 |
| *D. abbreviatus* | 97.90 | 1.80 | 99.70 | 653 | 1,623,139,935 | 32.17 | 7,800,247 | 1,733,013 | 52 | 222 | 85.12 | 6.90 | 92.02 |
| *D. ponderosae* | 91.96 | 5.70 | 97.66 | 2,112 | 223,596,823 | 35.82 | 16,553,750 | 3,217,956 | 4 | 12 | 83.56 | 9.66 | 93.22 |
| *E. fornicatus* | 99.28 | 0.48 | 99.76 | 185 | 219,832,364 | 36.46 | 3,845,785 | 701,585 | 20 | 71 | 94.18 | 5.22 | 99.40 |
| *E. similis* | 99.34 | 0.36 | 99.70 | 128 | 177,600,458 | 36.08 | 5,137,944 | 2,020,283 | 13 | 34 | 93.88 | 5.64 | 99.52 |
| *H. hampei* | 90.16 | 4.68 | 94.84 | 8,198 | 162,571,498 | 32.32 | 340,248 | 6,196 | 89 | 1,948 | 81.64 | 9.06 | 90.70 |
| *H. nitidus* | 90.34 | 6.18 | 96.52 | 210 | 231,406,857 | 35.19 | 16,438,241 | 7,158,500 | 6 | 14 | 81.34 | 11.40 | 92.74 |
| *I. typographus* | 94.06 | 5.70 | 99.76 | 272 | 236,816,287 | 35.21 | 6,654,004 | 438,776 | 12 | 74 | 86.26 | 10.50 | 96.76 |
| *P. cervinus* | 99.04 | 0.54 | 99.58 | 15 | 713,374,358 | 32.77 | 72,832,772 | 43,617,443 | 4 | 9 | 87.88 | 5.70 | 93.58 |
| *P. cylindrus* | 97.06 | 0.72 | 97.78 | 26 | 147,483,955 | 31.06 | 15,234,454 | 11,925,517 | 3 | 7 | 90.16 | 6.60 | 96.76 |
| *R. ferrugineus* | 96.04 | 1.98 | 98.02 | 24,005 | 589,402,552 | 32.21 | 471,583 | 7,985 | 268 | 5,093 | 88.24 | 7.26 | 95.50 |
| *S. oryzae* | 97.66 | 1.44 | 99.10 | 2,025 | 770,567,808 | 32.63 | 2,860,826 | 544,738 | 73 | 298 | 88.96 | 6.60 | 95.56 |
| *T. bicolor* | 97.90 | 1.56 | 99.46 | 517 | 575,240,612 | 36.23 | 48,296,002 | 43,417,794 | 5 | 10 | 87.28 | 6.00 | 93.28 |
| *T. castaneum* | 99.58 | 0.36 | 99.94 | 149 | 241,861,439 | 31.45 | 20,636,470 | 4,896,840 | 5 | 11 | 93.70 | 5.46 | 99.16 |
| *X. affinis* | 98.56 | 0.36 | 98.92 | 92,424 | 219,084,373 | 38.32 | 4,846,381 | 622 | 12 | 52,821 | 89.74 | 4.98 | 94.72 |
| *X. bispinatus* | 98.80 | 0.24 | 99.04 | 22,364 | 156,953,107 | 37.05 | 8,207,581 | 14,224 | 7 | 665 | 90.52 | 4.50 | 95.02 |
| *X. germanus* | 99.22 | 0.42 | 99.64 | 54 | 201,019,303 | 37.01 | 16,280,965 | 7,161,202 | 5 | 12 | 92.80 | 4.92 | 97.72 |
| *X. glabratus* | 97.18 | 1.14 | 98.32 | 101,578 | 216,324,869 | 39.65 | 125,395 | 6,336 | 24 | 5037 | 56.09 | 39.77 | 95.86 |
| *X. saxesenii* | 95.74 | 0.30 | 96.04 | 63 | 167,473,680 | 37.03 | 10,839,340 | 6,304,644 | 7 | 15 | 89.50 | 4.68 | 94.18 |

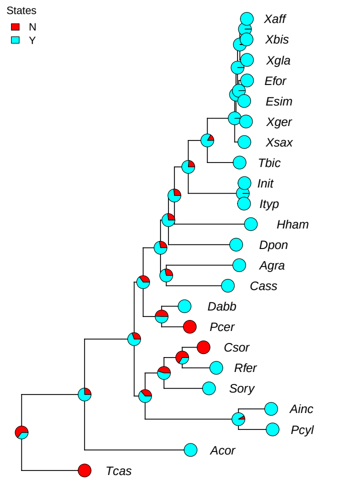


(a) Nest building

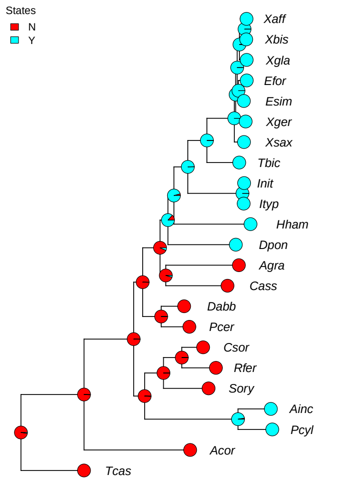


(b) Egg attendance

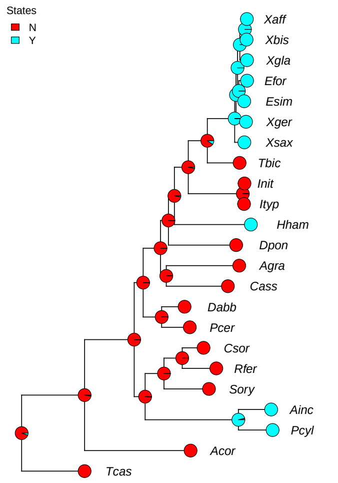


(c) Larval attendance

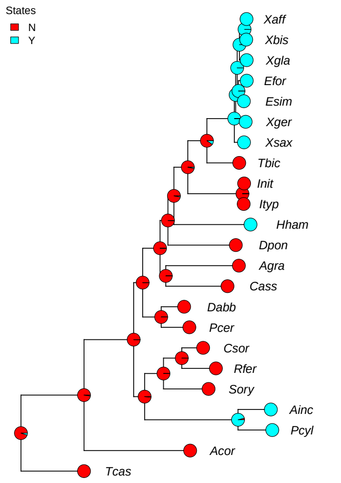


(d) Overlapping generations

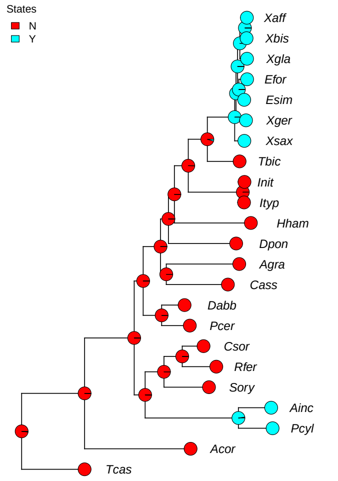


(e) Alloparental care

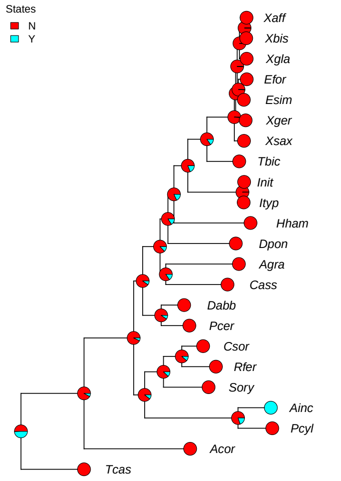


(f) Sterile workers

Figure S3: Social traits were reconstructed using ancestral reconstruction with phytools. State (Y/N) = social trait present/absent. Species abbreviations: Acor = *A. coryli*, Agra = *A. grandis grandis*, Ainc = *A. incompertus*, Cass = *C. assimilis*, Csor = *C. sordidus*, Dabb = *D. abbreviatus*, Dpon = *D. ponderosae*, Efor = *E. fornicatus*, Esim = *E. similis*, Hham = *H. hampei*, Init = *I. nitidus*, Ityp = *I. typographus*, Pcer = *P. cervinus*, Pcyl = *P. cylindrus*, Rfer = *R. ferrugineus*, Sory = *S. oryzae*, Tbic = *T. bicolor*, Tcas = *T. castaneum*, Xaff = *X. affinis*, Xbis = *X. bispinatus*, Xger = *X. germanus*, Xgla = *X. glabratus*, Xsax = *X. saxesenii*.

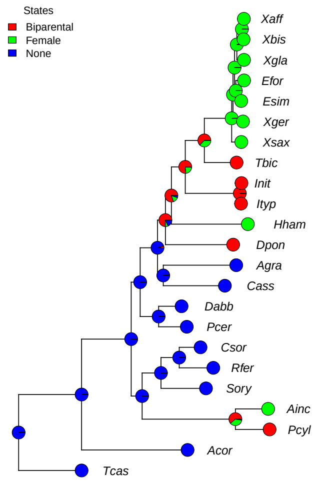


(a) Care pattern

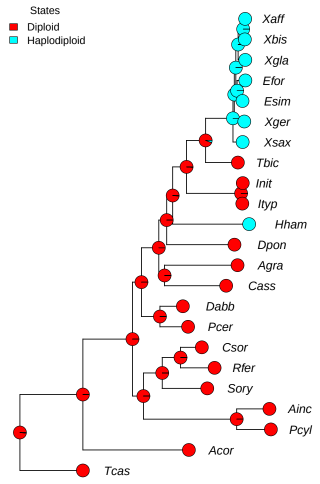


(b) Sex determination system

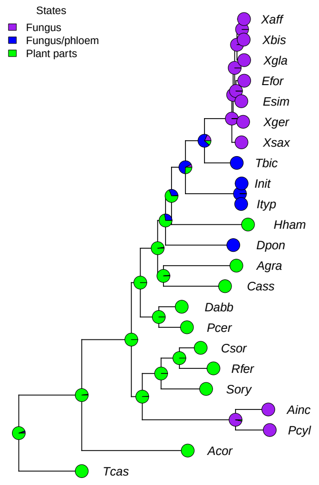


(c) Nutrition

Figure S4: Traits were reconstructed using ancestral reconstruction with phytools. Species abbreviations: Acor = *A. coryli*, Agra = *A. grandis grandis*, Ainc = *A. incompertus*, Cass = *C. assimilis*, Csor = *C. sordidus*, Dabb = *D. abbreviatus*, Dpon = *D. ponderosae*, Efor = *E. fornicatus*, Esim = *E. similis*, Hham = *H. hampei*, Init = *I. nitidus*, Ityp = *I. typographus*, Pcer = *P. cervinus*, Pcyl = *P. cylindrus*, Rfer = *R. ferrugineus*, Sory = *S. oryzae*, Tbic = *T. bicolor*, Tcas = *T. castaneum*, Xaff = *X. affinis*, Xbis = *X. bispinatus*, Xger = *X. germanus*, Xgla = *X. glabratus*, Xsax = *X. saxesenii.*

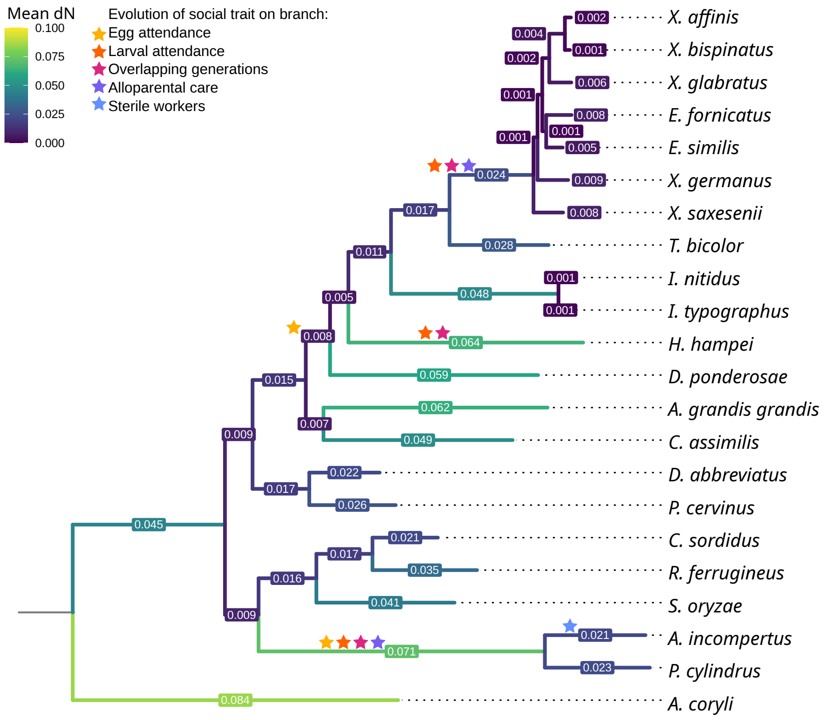


Figure S5: Mean dN for each branch across the phylogenetic tree.

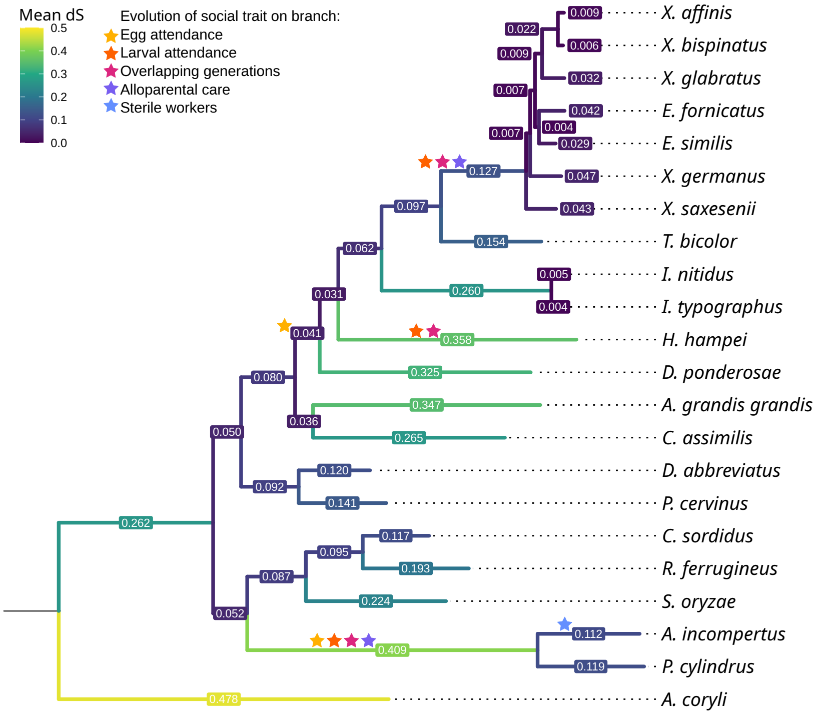


Figure S6: Mean dS for each branch across the phylogenetic tree.

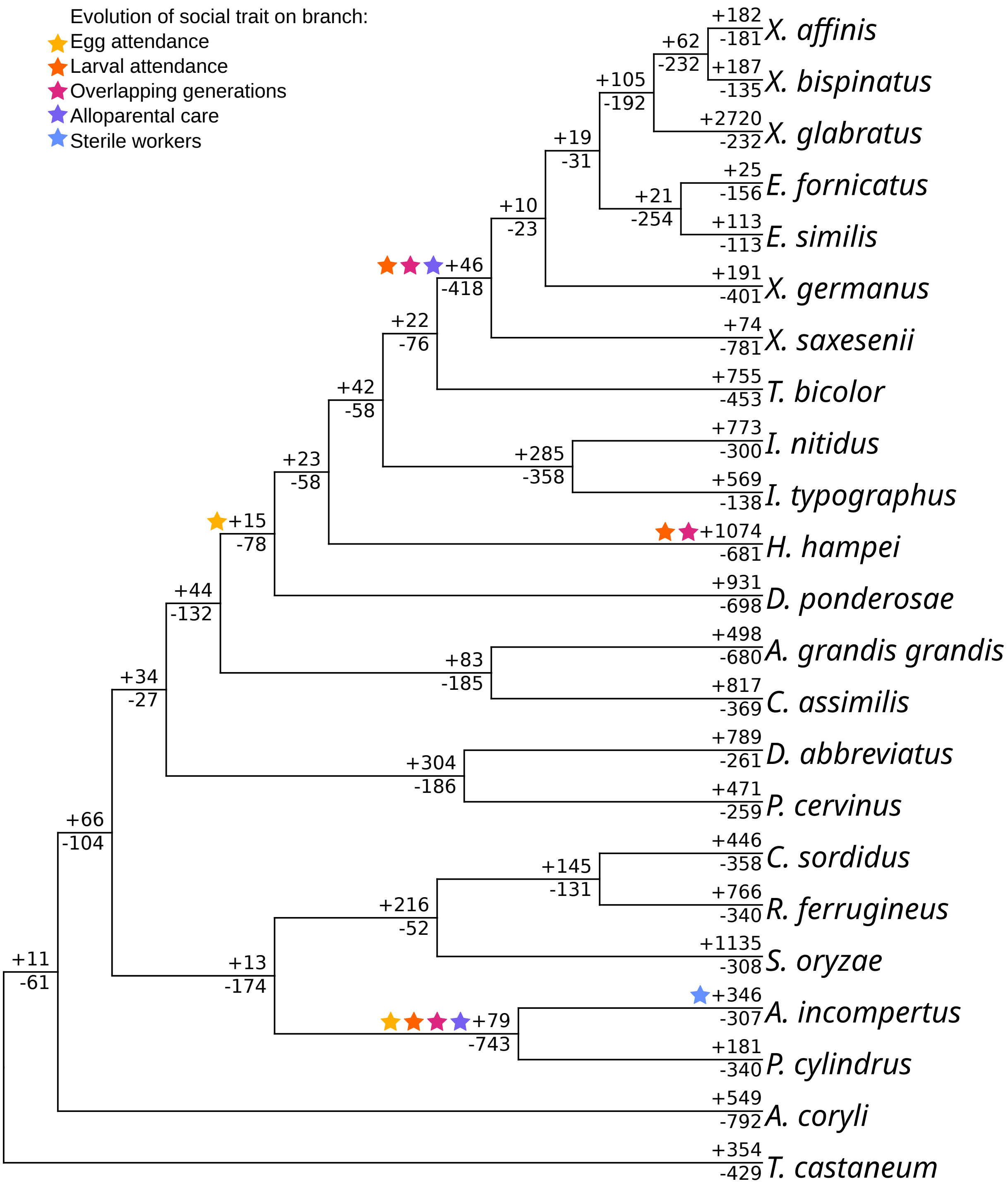


Figure S7: Phylogenetic tree showing number of expansions (+) and contractions (-) of gene families for each branch. Only gene families present at root are analysed. Stars denote the evolution of social traits according to the legend.

Table S3: Numbers and functional enrichment of expansions and contractions of gene families on branches of interest. Sig. changes indicates the number of significantly expanded and contracted genes per branch. Functional enrichment was analysed using GO-term enrichment analysis. All GO terms with $p<0.05$ are shown in order of decreasing significance. Bold font indicates FDR significance.

| Branch | Sig. changes | Functional enrichment |
|---|---|---|
| Expansions | | |
| *A. incompertus* | 120 | - Microtubule-based process |
| *H. hampei* | 280 | - Signal transduction<br>- Cell communication<br>- Cellular response to stimulus<br>- Response to stimulus<br>- Signaling<br>- Phosphorus metabolic process |
| Platypodinae ancestor | 19 | No Enriched GO terms |
| Scolytinae ancestor | 12 | No Enriched GO terms |
| Xyleborinae ancestor | 29 | No Enriched GO terms |
| Contractions | | |
| *A. incompertus* | 113 | **- Response to stress**<br>**- DNA metabolic process**<br>- Regulation of molecular function<br>- Multicellular organismal process<br>- Cellular response to stress<br>- DNA repair |
| *H. hampei* | 42 | - Proteolysis<br>- Protein metabolic process<br>- Aromatic compound biosynthetic process<br>- Organic cyclic compound biosynthetic process<br>- Heterocycle biosynthetic process |
| Platypodinae ancestor | 80 | **- Multicellular organismal process**<br>- Transmembrane transport<br>- Proteolysis |
| Scolytinae ancestor | 46 | - DNA replication<br>- Proteolysis |
| Xyleborinae ancestor | 101 | **- Proteolysis**<br>- Multicellular organismal process<br>- Cellular aromatic compound metabolic process<br>- Heterocycle metabolic process<br>- DNA metabolic process |

Table S4: Numbers and functional annotation of expansions and contractions of gene families exclusively present on one branch of interest (not present on any other branch within the dataset).

| Branch | Number | Functional annotation |
|---|---|---|
| Expansions | | |
| *A. incompertus* | 7 | - Structural constituent of ribosome \| Ribosomal L37ae protein family |
| *H. hampei* | 21 | - Belongs to the cytochrome b5 family \| Cytochrome b5<br>- Mediates the side-chain deamidation of N-terminal glutamine residues to glutamate, an important step in N-end rule pathway of protein degradation<br>- Protein-kinase domain of FAM69<br>- Transmembrane protein 188<br>- Trypsin-like serine protease |
| Platypodinae ancestor | 0 | NA |
| Scolytinae ancestor | 0 | NA |
| Xyleborini ancestor | 1 | NA |
| Contractions | | |
| *A. incompertus* | 24 | - C-type lectin (CTL) or carbohydrate-recognition domain (CRD) actin filament binding \| Apx/Shroom domain ASD2<br>- Calcium ion binding<br>- Cation efflux family<br>- Mab-21 \| Leucine rich repeat<br>- NHL repeat \| Zinc ion binding \| Tetraspanin family |
| *H. hampei* | 4 | - Spaetzle |
| Platypodinae ancestor | 7 | - Catalyzes the reduction of fatty acyl-CoA to fatty alcohols electron carrier activity<br>- Insect cuticle protein \| Structural constituent of cuticle<br>- Major Facilitator Superfamily<br>- Belongs to the type-B carboxylesterase lipase family \| Carboxylesterase family |
| Scolytinae ancestor | 21 | - heme binding \| Cytochrome P450<br>- Belongs to the adenylyl cyclase class-4 guanylyl cyclase family<br>- Cognition |

| | | |
|---|---|---|
| | | - Cell-cell signaling by wnt<br>- Chitin binding |
| Xyleborini ancestor | 40 | - Carbohydrate binding<br>- Glyceraldehyde 3-phosphate dehydrogenase, NAD binding domain \| oxidoreductase activity, acting on the aldehyde or oxo group of donors, NAD or NADP as acceptor<br>- Belongs to the glycosyltransferase 11 family<br>- Adenylate cyclase type 10-like \| Adenylate and Guanylate cyclase catalytic domain<br>- Cyclic nucleotide-monophosphate binding domain \| Hyperpolarization activated cyclic nucleotide-gated potassium channel<br>- SERine Proteinase INhibitors \| Belongs to the serpin family<br>- Transglutaminase-like superfamily \| F-box only protein 21<br>- Belongs to the TRAFAC class myosin-kinesin ATPase superfamily. Kinesin family \| ATP-dependent microtubule motor activity, plus-end-directed<br>- DNA binding<br>- NAD(P)H oxidase activity<br>- Receptor activity<br>- Zinc finger protein 142<br>- Zinc ion binding<br>- Axonemal central apparatus assembly |

Table S5: Gene functions of convergently contracted or expanded gene families on different branches of interest. The overlaps correspond to overlaps between expansions or contractions seen in the SuperExactTest. Overlaps found in Figure 4 but not listed here had no assigned functions. Letters in superscript for some functions indicate which high-level functions are assigned in Table 2.

| **Overlap** | **Gene function** |
|---|---|
| **Contractions** | |
| Xyleborini ancestor &<br><br>*A. incompertus* | - Belongs to the peptidase C1 family \| Cysteine-type peptidase activity<br>- Belongs to the glycosyl hydrolase 1 family \| Glycosyl hydrolase family 1 |

| | |
|---|---|
| | - Belongs to the type-B carboxylesterase lipase family \| carboxylic ester hydrolase activity<br>- Catalyzes juvenile hormone hydrolysis<br>- Prostaglandin reductase<br>- Enoyl-(Acyl carrier protein) reductase<br>- Aldo/keto reductase family \| Oxidoreductase activity<br>- Phospholipid scrambling \| Protein phosphatase 1 regulatory subunit 15A<br>- Belongs to the major facilitator superfamily. Sugar transporter (TC 2.A.1.1) family \| glucose import<br>- Anterograde trans-synaptic signaling |
| Xyleborini ancestor & Platypodinae ancestor | - Myb/SANT-like DNA-binding domain [a]<br>- Transferase activity, transferring hexosyl groups [b]<br>- Belongs to the glycosyl hydrolase 1 family \| Glycosyl hydrolase family 1 [b]<br>- Belongs to the type-B carboxylesterase lipase family [c]<br>- Belongs to the type-B carboxylesterase lipase family \| carboxylic ester hydrolase activity [c]<br>- Choline dehydrogenase activity \| GMC oxidoreductase \| Belongs to the GMC oxidoreductase family [d]<br>- Heat shock protein \| Heat shock 70 kDa protein cognate [d]<br>- Prostaglandin reductase [e]<br>- Aldo/keto reductase family \| Oxidoreductase activity [e]<br>- Macromolecule localization \| transporter activity [f]<br>- Trypsin-like serine protease \| serine-type endopeptidase activity \| Belongs to the peptidase S1 family [i]<br>- Thrombospondin type 1 domain [i] |
| Platypodinae ancestor & *H. hampei* | - DNA binding \| nucleosome assembly [a]<br>- Protein heterodimerization activity \| C-terminus of histone H2A \| Histone H2A [a]<br>- Calcium binding and coiled-coil domain (CALCOCO1) like [a]<br>- Belongs to the type-B carboxylesterase lipase family [c]<br>- Heat shock protein \| Heat shock 70 kDa protein cognate [d]<br>- Choline dehydrogenase activity \| GMC oxidoreductase \| Belongs to the GMC oxidoreductase family [d]<br>- Glutathione s-transferase \| Glutathione S-transferase, C-terminal domain [e]<br>- Enoylreductase [g]<br>- Protein tyrosine phosphatase activity[i] |

| | |
|---|---|
| | - Trypsin-like serine protease \| serine-type endopeptidase activity \| Belongs to the peptidase S1 family [i]<br>- Peptidase family M1 domain [i]<br>- Thrombospondin type 1 domain [i] |
| Xyleborini ancestor &<br>*H. hampei* | - Belongs to the type-B carboxylesterase lipase family [c]<br>- Heat shock protein \| Heat shock 70 kDa protein cognate [d]<br>- Choline dehydrogenase activity \| GMC oxidoreductase \| Belongs to the GMC oxidoreductase family [d]<br>- Heme binding \| Cytochrome P450 [e]<br>- Glycosyl hydrolase family 79, N-terminal domain [e]<br>- Belongs to the short-chain dehydrogenases reductases (SDR) family \| retinol dehydrogenase [i]<br>- Trypsin-like serine protease \| serine-type endopeptidase activity \| Belongs to the peptidase S1 family [i]<br>- Belongs to the peptidase C1 family \| Cysteine-type peptidase activity [i]<br>- Thrombospondin type 1 domain [i] |
| Scolytinae ancestor & Platypodinae ancestor | - Belongs to the small heat shock protein (HSP20) family \| Hsp20/alpha crystallin family [d]<br>- Belongs to the CRISP family [i]<br>- Ionotropic glutamate receptor activity [h] |
| *A. incompertus* &<br>*H. hampei* | - nucleosomal DNA binding \| Histone H3 \| Histone H3.3-like<br>- Belongs to the short-chain dehydrogenases reductases (SDR) family<br>- Belongs to the peptidase C1 family \| Cysteine-type peptidase activity<br>- Histone H2B \| Protein heterodimerization activity<br>- Core component of nucleosome |
| Xyleborini ancestor &<br><br>Scolytinae ancestor | - Cyclic nucleotide-monophosphate binding domain \| Hyperpolarization activated cyclic nucleotide-gated potassium channel<br>- Phospholipid scrambling \| Protein phosphatase 1 regulatory subunit 15A<br>- Belongs to the major facilitator superfamily. Sugar transporter (TC 2.A.1.1) family \| glucose import |
| Platypodinae ancestor &<br><br>*A. incompertus* | - Belongs to the glycosyl hydrolase 1 family \| Glycosyl hydrolase family 1<br>- Belongs to the type-B carboxylesterase lipase family \| carboxylic ester hydrolase activity<br>- Prostaglandin reductase<br>- Aldo/keto reductase family \| Oxidoreductase activity |

| | |
|---|---|
| Xyleborini ancestor & Platypodinae ancestor & *A. incompertus* | - Belongs to the glycosyl hydrolase 1 family \| Glycosyl hydrolase family 1<br>- Belongs to the type-B carboxylesterase lipase family \| carboxylic ester hydrolase activity<br>- Prostaglandin reductase<br>- Aldo/keto reductase family \| Oxidoreductase activity |
| Xyleborini ancestor & Platypodinae ancestor & *H. hampei* | - Belongs to the type-B carboxylesterase lipase family [c]<br>- Heat shock protein \| Heat shock 70 kDa protein cognate [d]<br>- Choline dehydrogenase activity \| GMC oxidoreductase \| Belongs to the GMC oxidoreductase family [d]<br>- Trypsin-like serine protease \| serine-type endopeptidase activity \| Belongs to the peptidase S1 family[i]<br>- Thrombospondin type 1 domain [i] |
| Scolytinae ancestor & *A. incompertus* | - Phospholipid scrambling \| Protein phosphatase 1 regulatory subunit 15A<br>- Belongs to the major facilitator superfamily. Sugar transporter (TC 2.A.1.1) family \| glucose import |
| *A. incompertus* & Scolytinae ancestor & Xyleborini ancestor | - Phospholipid scrambling \| Protein phosphatase 1 regulatory subunit 15A<br>- Belongs to the major facilitator superfamily. Sugar transporter (TC 2.A.1.1) family \| Glucose import |
| *A. incompertus* & *H. hampei* Xyleborini ancestor | - Belongs to the peptidase C1 family \| Cysteine-type peptidase activity |
| **Expansions** | |
| *A. incompertus* & *H. hampei* | - Facilitated trehalose transporter Tret1-2 homolog-like Protein<br>- large 1 tumor suppressor \| Guanylate kinase homologues<br>- Adenylate cyclase type 10-like \| Adenylate and Guanylate cyclase catalytic domain<br>- Belongs to the TRAFAC class myosin-kinesin ATPase superfamily. Kinesin family \| ATP-dependent microtubule motor activity, plus-end-directed |
| Platypodinae ancestor & *H. hampei* | - Cytochrome c oxidase subunit Vb \| Cytochrome-c oxidase activity [e] |

Table S6: Species specific genes in *A. incompertus*. All genes/orthogroups exclusively present in *A. incompertus* according to OrthoFinder analysis. Genes were functionally annotated using eggnogmapper.

| Trait | Functional annotation |
|---|---|
| Social Biology / Reproductive Regulation | - Juvenile hormone binding protein domains in insects<br>- Component of the sequence-specific heterotrimeric transcription factor (NF-Y) which specifically recognizes a 5'- CCAAT-3' box motif found in the promoters of its target genes. NF- Y can function as both an activator and a repressor, depending on its interacting cofactors<br>- Component of the sequence-specific heterotrimeric transcription factor (NF-Y) which specifically recognizes a 5'- CCAAT-3' box motif found in the promoters of its target genes. NF- Y can function as both an activator and a repressor, depending on its interacting cofactors<br>- Homeodomain<br>- Histone H1<br>- Histone acetylation protein |
| Signaling | - Protein serine threonine kinase activity<br>- Protein serine threonine kinase activity<br>- Protein tyrosine kinase<br>- Phosphatidylinositol phospholipase C activity<br>- Belongs to the protein-tyrosine phosphatase family. Non-receptor class myotubularin subfamily<br>- Guanylate kinase homologues |
| Host-Plant Adaptation | - Cytochrome P450<br>- Trehalase<br>- Oxidoreductase<br>- Enoyl-(Acyl carrier protein) reductase<br>- Aspartate oxidase activity |
| Protein Turnover | - SCF-dependent proteasomal ubiquitin-dependent protein catabolic process \| F-box domain<br>- Ubiquitin homologues<br>- Ubiquitin-like domain<br>- Proteasome-substrate-size regulator, mid region |
| Vesicular Trafficking | - Vacuolar protein sorting-associated protein \| Vacuolar-sorting-associated 13 protein C-terminal<br>- Vacuolar protein sorting-associated protein<br>- Adaptor-related protein complex 2, mu 1 subunit<br>- Intraflagellar transport |

<table>
<tr><td>Gene Expression Regulation</td><td>- RNA splicing<br>- Pre-mRNA branch point binding<br>- Acts as an adapter for the XPO1 CRM1-mediated export of the 60S ribosomal subunit</td></tr>
<tr><td>Mitochondrial respiratory chain</td><td>- Mitochondrial carrier protein<br>- Arginine methyltransferase involved in the assembly or stability of mitochondrial NADH ubiquinone oxidoreductase complex (complex I)<br>- Belongs to the complex I 30 kDa subunit family</td></tr>
<tr><td>Other</td><td>- K02A2.6-like<br>- Acid phosphatase activity | Raptor N-terminal CASPase like domain | protein of MTOR<br>- NAD(P)-binding Rossmann-like domain<br>- Transmembrane Fragile-X-F protein<br>- Acetylgalactosaminyltransferase activity<br>- Cadherin repeats<br>- Fringe-like<br>- Metal ion binding<br>- Pyruvate kinase activity<br>- Domain of unknown function (DUF4485)<br>- Protein of unknown function (DUF3808)</td></tr>
</table>

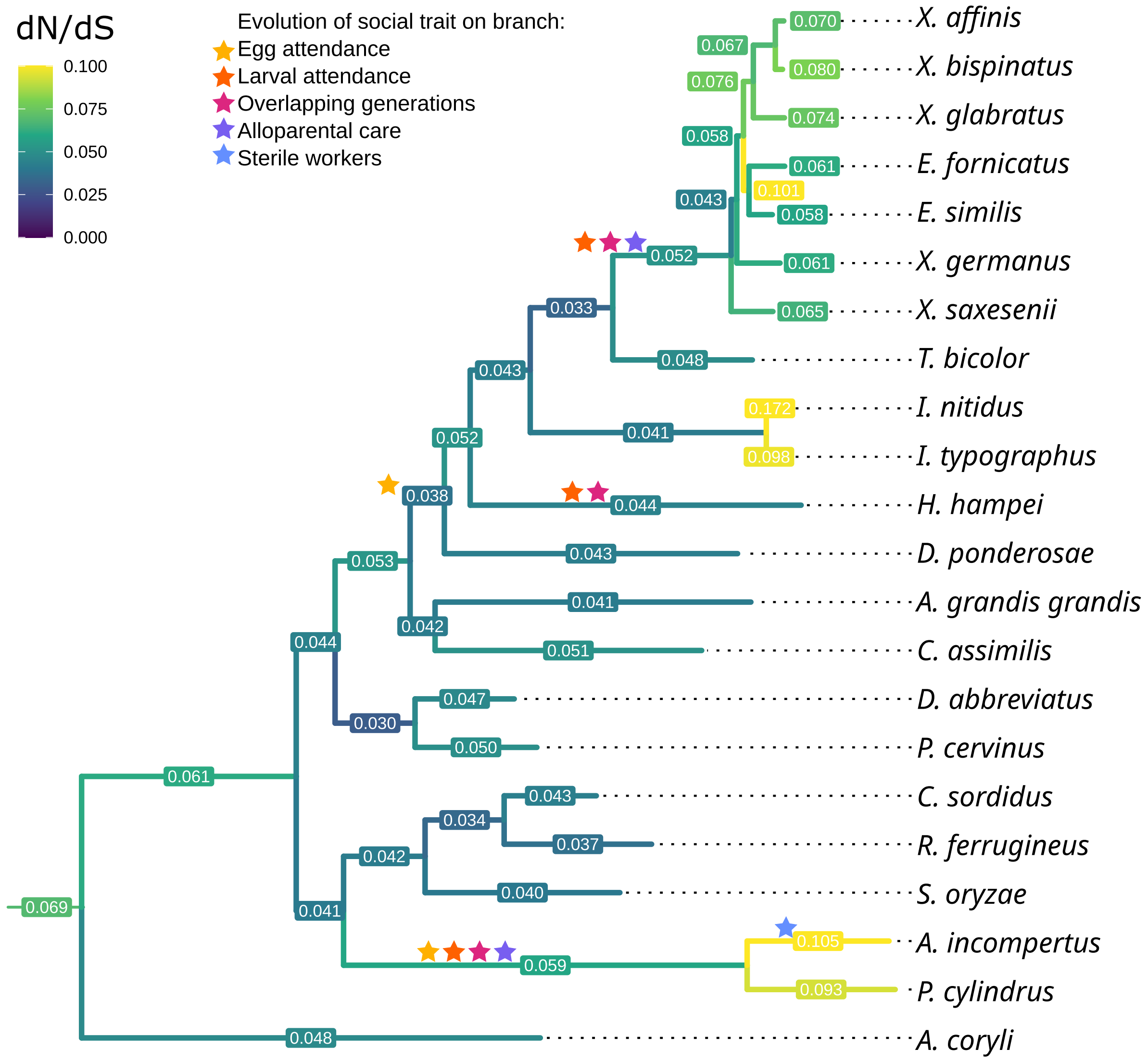


Figure S8: Phylogenetic tree showing concatenated branch dN/dS for all single copy orthologs.

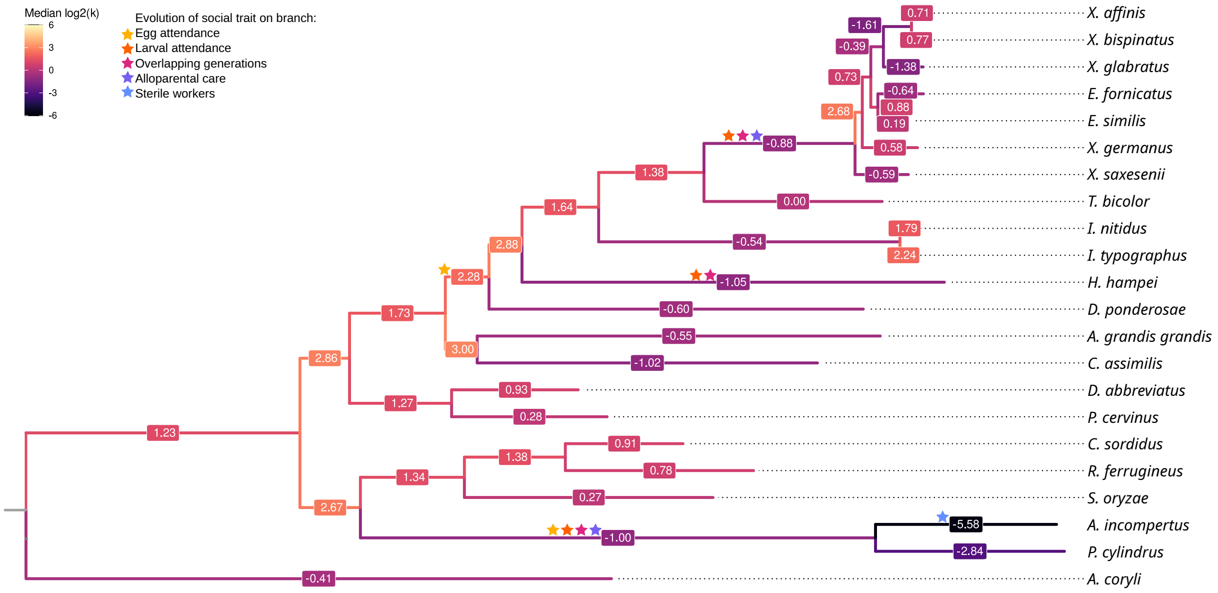


Figure S9: Phylogenetic tree showing median log2(k) per branch, where k is the relaxation and intensification parameter. Positive values indicate relaxed selection, while negative values indicate relaxation of selection.

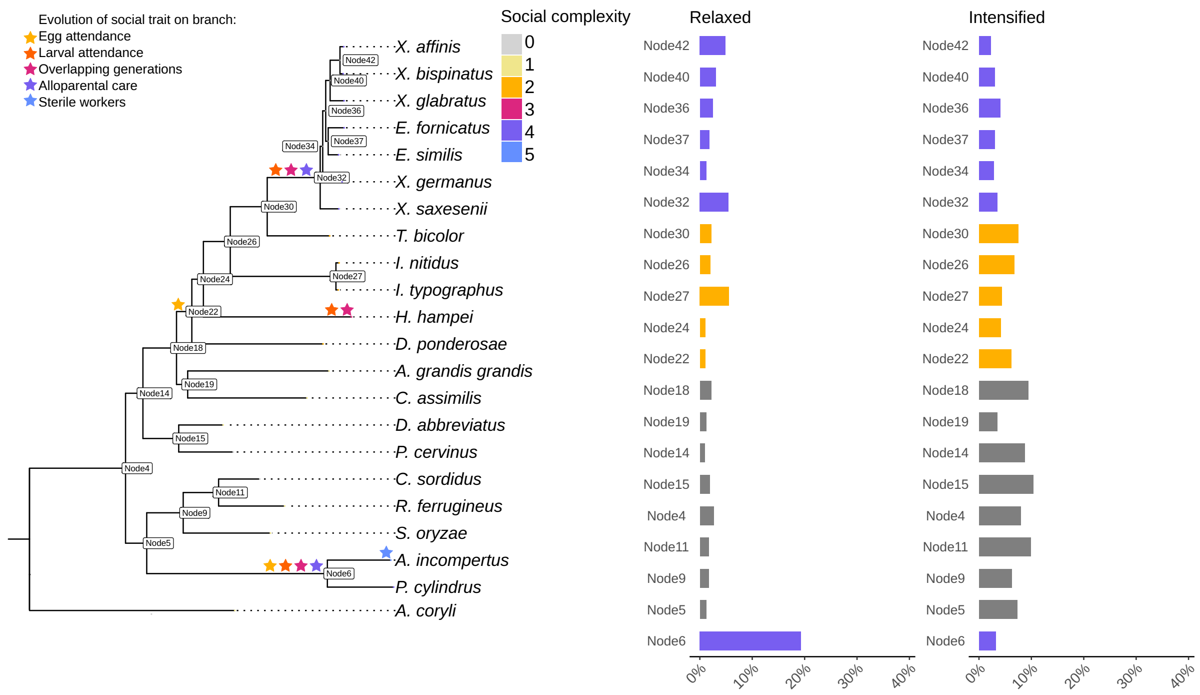


Figure S10: Proportion of single copy orthologues under relaxed or intensified selection for each internal node.

Table S7: PGLS model model comparison. Models tested: Brownian: $gls(median_k \sim Sociality,\ correlation = corBrownian(1, tree, form =\sim Species)$. OU: $gls(median_k \sim Sociality,\ correlation = corMartins(1, tree, form =\sim Species)$. df: degrees of freedom, AIC: Akaike information criterion, BIC: Bayesian information criterion, logLik: log Likelihood, L.Ratio: likelihood ratio.

| | Model | df | AIC | BIC | logLik | Test | L.Ratio | p-value |
|---|---|---|---|---|---|---|---|---|
| Brownian | 1 | 7 | -8.64 | -0.69 | 11.32 | | | |
| OU | 2 | 8 | -16.54 | -7.46 | 16.27 | 1 vs 2 | 9.90 | 0.0017 |

Table S8: Estimated marginal means for each sociality level in the PGLS with OU model. emmean: estimated marginal mean, SE: standard error, df: degrees of freedom, CL: confidence interval. Degrees-of-freedom method: satterthwaite, confidence level used: 0.95.

| Sociality level | emmean | SE | df | lower.CL | upper.CL |
|---|---|---|---|---|---|
| no | 1.039 | 0.0938 | 17.1 | 0.841 | 1.236 |
| nb | 1.034 | 0.0682 | 12.6 | 0.886 | 1.182 |
| ea | 0.872 | 0.0951 | 13.9 | 0.668 | 1.077 |
| og | 0.921 | 0.1620 | 18.6 | 0.582 | 1.260 |
| ac | 0.842 | 0.1010 | 12.4 | 0.622 | 1.062 |
| sw | 0.504 | 0.1610 | 20.0 | 0.169 | 0.839 |

Table S9: Contrasts between sociality level in the PGLS of the relaxation and intensification parameter (median k) with OU model. estimate: estimated difference in means, SE: standard error, df: degrees of freedom, CL: confidence interval, t.ratio: t statistic. Degrees-of-freedom method: satterthwaite, P value adjustment: FDR.

| Sociality level | Estimate | SE | df | t.ratio | p.value |
|---|---|---|---|---|---|
| no care - nest building | 0.00441 | 0.107 | 23.5 | 0.041 | 0.9674 |
| no care - egg attendance | 0.16614 | 0.132 | 18.7 | 1.260 | 0.3720 |
| no care - overlapping generations | 0.11743 | 0.186 | 19.7 | 0.631 | 0.7298 |
| no care - alloparental care | 0.19690 | 0.137 | 16.7 | 1.438 | 0.3306 |
| no care - sterile workers | 0.53439 | 0.185 | 20.4 | 2.882 | 0.0683 |
| nest building - egg attendance | 0.16174 | 0.115 | 17.7 | 1.408 | 0.3306 |
| nest building - overlapping generations | 0.11303 | 0.174 | 19.4 | 0.648 | 0.7298 |
| nest building - alloparental care | 0.19249 | 0.121 | 15.0 | 1.594 | 0.3294 |
| nest building - sterile workers | 0.52998 | 0.174 | 19.9 | 3.046 | 0.0683 |
| egg attendance - overlapping generations | -0.04871 | 0.185 | 20.4 | -0.263 | 0.8754 |
| egg attendance - alloparental care | 0.03076 | 0.131 | 20.6 | 0.234 | 0.8754 |
| egg attendance - sterile workers | 0.36825 | 0.186 | 20.0 | 1.983 | 0.2463 |
| overlapping generations - alloparental care | 0.07947 | 0.189 | 18.7 | 0.419 | 0.8497 |
| overlapping generations - sterile workers | 0.41696 | 0.228 | 19.9 | 1.831 | 0.2463 |
| alloparental care - sterile workers | 0.33749 | 0.181 | 23.2 | 1.862 | 0.2463 |

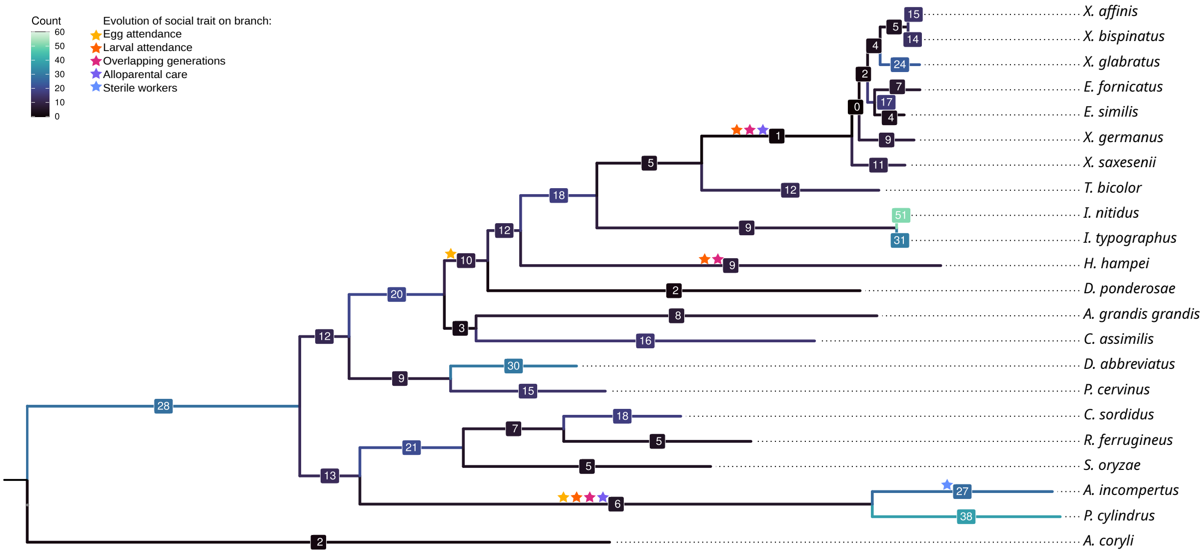


Figure S11: Phylogenetic tree showing the number of genes under positive selection on each branch.

Table S10: Gene IDs and full gene names in *T. castaneum* for all genes under positive selection on the indicated branch.

| | |
|---|---|
| *A. incompertus* | |
| noc | zinc finger protein no ocelli |
| LOC663869 | longitudinals lacking protein |
| nej | CREB binding protein nejire |
| E(Pc) | Enhancer of Polycomb |
| gcm | glial cells missing |
| LOC655000 | solute carrier family 35 member G1 |
| LOC655136 | Hermansky-Pudlak syndrome 3 protein homolog |
| LOC655185 | putative sodium-coupled neutral amino acid transporter 10 |
| NaPi-III | Na[+]-dependent inorganic phosphate cotransporter type III |
| LOC654928 | escargot |
| Hers | Histone gene-specific Epigenetic Repressor in late S phase |
| br | broad-complex core protein |
| Msh | muscle segment homeodomain protein |
| LOC657981 | kinesin 13C |
| LOC657816 | leucine-rich repeat-containing protein 49 |
| cmb | combover |
| mei-9 | DNA repair endonuclease XPF mei-9 |
| Nt5c | 5' nucleotidase C |
| Usp15-31 | Ubiquitin specific protease 15/31 |
| LOC657696 | skin secretory protein xP2 |
| stv | BAG domain-containing protein starvin |
| pcx | pecanex |
| LOC664505 | adipocyte plasma membrane-associated protein Hemomucin |
| LOC655102 | uncharacterized LOC655102 |
| LOC663924 | hypothetical protein |
| LOC664527 | hypothetical protein |
| LOC103314814 | uncharacterized LOC103314814 |
| *H. hampei* | |
| slbo | slow border cells |
| Asx | transcriptional regulator additional sex combs |
| LOC659686 | protein Asterix |
| LOC660343 | protein tramtrack |
| Ranbp16 | Ran-binding protein 16 |
| LOC664496 | anoctamin-1 |
| LOC661834 | myocardin-related transcription factor B |
| LOC100142298 | uncharacterized LOC100142298 |
| LOC103314293 | uncharacterized LOC103314293 |
| Platypodinae ancestor | |
| LOC661994 | U2 small nuclear ribonucleoprotein auxiliary factor 35 kDa subunit-related protein 2 |
| LOC661424 | deleted in azoospermia protein 2 |
| LOC107397896 | titin-like |
| nej | CREB binding protein nejire |
| LOC659686 | protein Asterix |
| ND-ACP | NADH dehydrogenase (ubiquinone) acyl carrier protein |
| Scolytinae ancestor | |
| LOC655055 | hemicentin-2 |
| br | broad-complex core protein |
| LOC655175 | coiled-coil domain-containing protein 130 homolog |
| Sarnp | SAP domain containing ribonucleoprotein |
| Ino80 | chromatin-remodeling ATPase INO80 |
| LOC661190 | KN motif and ankyrin repeat domain-containing protein 2 |
| LOC659765 | diacylglycerol kinase theta |
| LOC662169 | cyclin-dependent kinase-like 4 |
| Chchd3 | Coiled-coil-helix-coiled-coil-helix domain containing 3 |
| LOC661642 | tetratricopeptide repeat protein 28 |
| Xyleborini ancestor | |
| LOC103312803 | early growth response protein 1 |
